# Translation re-initiation after uORFs does not fully protect mRNAs from nonsense-mediated decay

**DOI:** 10.1101/2022.01.10.475702

**Authors:** Paul J. Russell, Jacob A. Slivka, Elaina P. Boyle, Arthur H. M. Burghes, Michael G. Kearse

## Abstract

It is estimated that nearly 50% of mammalian transcripts contain at least one upstream open reading frame (uORF), which are typically one to two orders of magnitude smaller than the downstream main ORF. Most uORFs are thought to be inhibitory as they sequester the scanning ribosome, but in some cases allow for translation re-initiation. However, termination in the 5ʹ UTR at the end of uORFs resembles pre-mature termination that is normally sensed by the nonsense-mediated mRNA decay (NMD) pathway. Translation re-initiation has been proposed as a method for mRNAs to prevent NMD. Here we test how uORF length influences translation re-initiation and mRNA stability in HeLa cells. Using custom 5ʹ UTRs and uORF sequences, we show that re-initiation can occur on heterologous mRNA sequences, favors small uORFs, and is supported when initiation occurs with more initiation factors. After determining reporter mRNA half-lives in HeLa cells and mining available mRNA half-life datasets for cumulative predicted uORF length, we conclude that translation re-initiation after uORFs is not a robust method for mRNAs to prevent NMD. Together, these data suggests that the decision of whether NMD ensues after translating uORFs occurs before re-initiation in mammalian cells.

## INTRODUCTION

Canonical eukaryotic translation initiation follows a cap- and scanning-dependent mode (reviewed extensively in Hinnebusch 2017; Shirokikh and Preiss 2018). First, the 5ʹ m^7^G cap recruits the eIF4F complex via the eIF4E cap-binding protein. eIF4F also provides an opportunity for the mRNA to form a close-loop conformation with the poly(A) tail and PABP that is thought to increase translation efficiency. eIF4F then recruits the 43S pre-initiation complex (PIC), which is comprised of the 40S small ribosomal subunit, eIF1, eIF1A, eIF3, eIF5, and the ternary complex (eIF2•GTP•Met-tRNAi^Met^). The 43S PIC then scans 5ʹ to 3ʹ in search of an AUG start codon (Brito Querido et al. 2020; Wang et al. 2022). The 48S initiation complex is formed after start codon recognition and eIF2 subsequently hydrolyzes GTP to release Met-tRNAi^Met^.

After initiation factors dissociate, initiation is completed once the 60S large ribosomal subunit joins with aid from eIF5B•GTP to form the complete 80S ribosome (Lapointe et al. 2022).

Due to the 5ʹ to 3ʹ nature of the scanning 43S PIC, the first AUG start codon in the mRNA is often recognized most efficiently and is primarily used for protein synthesis—albeit the surrounding context (i.e., Kozak sequence) and distance from the 5ʹ end influences start codon recognition (Kozak 1986, 1991, 1997; Brito Querido et al. 2020). A major hurdle for the ribosome from recognizing the AUG start codon of the protein encoding main open reading frame (ORF) is the presence of upstream start codons and upstream open reading frames (uORFs) in the 5ʹ untranslated region (UTR) (Young and Wek 2016; Gunisova et al. 2018).

Nearly 50% of mammalian 5ʹ UTRs harbor at least one uORF (Calvo et al. 2009). Some uORFs do not present much of an obstacle as recent work has shown that the footprint of the eIF4F complex creates a “blind spot” in the first ∼50 nucleotides (nts) for the ribosome and only AUG start codons after this point are efficiently recognized (Brito Querido et al. 2020). However, 5ʹ UTRs are often larger than the length of this blind spot and contain multiple uORFs. In fact, stress response mRNAs contain multiple evolutionarily conserved uORFs that are thought to be key regulators of translation (Young and Wek 2016; Wek 2018). For example, the *ATF4* mRNA 5ʹ UTR contains at least two uORFs (Lu et al. 2004; Vattem and Wek 2004; Pavitt and Ron 2012). It has been proposed for *ATF4* mRNA that most scanning PICs initiate at the start codon of uORF1 and that translation of uORF1 favors translation re-initiation at uORF2 under normal physiological conditions. Re-initiation occurs when a ribosome terminates and releases the polypeptide but fails to have the small 40S subunit recycled off and subsequently continues to scan downstream in search of a start codon (Vattem and Wek 2004; Young and Wek 2016).

Since uORF2 overlaps the ATF4 coding sequence and ends downstream from the beginning of the ATF4 coding sequence, ATF4 protein is not efficiently synthesized unless uORF2 is bypassed during cell stress (Vattem and Wek 2004; Young and Wek 2016; Wek 2018).

It has been proposed that re-initiation could negate the ribosome from triggering nonsense-mediated mRNA decay (NMD) during termination at uORFs in the 5ʹ UTR or at premature termination codons within the major ORF since the late steps of termination and/or ribosome recycling are not completed (Buisson et al. 2006; Paulsen et al. 2006; Stump et al. 2012; Hulsebos et al. 2014; Pereira et al. 2015; Jagannathan and Bradley 2016; Dyle et al. 2020). In mammalian cells, NMD is triggered by termination ≥50 nts upstream of an exon-junction complex (EJC) or upstream of long 3ʹ UTRs (Lykke-Andersen and Jensen 2015; Yi et al. 2021). Ribosomes that re-initiate would also displace EJCs and mRNA-bound Upf1 further preventing NMD during subsequent rounds of translation (Dyle et al. 2020). In this report, we test how uORF length affects translation re-initiation and mRNA stability. Our data show that translation re-initiation is favored after small uORFs but does not robustly prevent NMD. Reporter mRNAs with small or large uORFs that elicit re-initiation 100-fold differently have similar drastically shortened half-lives. Mining published mRNA half-life datasets for cumulative computationally predicted uORF length demonstrates that mRNAs with predicted uORFs of any length are equally less stable on average than mRNAs without predicted uORFs. These data suggests that the decision of whether NMD ensues from translating uORFs occurs before re-initiation in mammalian cells.

## RESULTS AND DISCUSSION

To specifically measure re-initiation and how it impacts mRNA stability, we designed nanoLuciferase (nLuc) reporters to harbor an uORF that maximally prevents leaky scanning of the PIC and avoids reporter signal from ribosome readthrough. This was achieved by using a synthetic 5ʹ UTR with three important elements: i) a 72 nt CAA-repeat leader sequence to allow for an unstructured sequence that is in the optimal length window for cap- and scanning-dependent initiation (Brito Querido et al. 2020), ii) an uORF comprised of three AUG start codons in perfect Kozak context (3XAUG), iii) and a 16 nt unstructured CAA-repeat linker between the uORF and nLuc start codon to separate and frameshift the ORFs (**Fig. 1A**).

**Figure 1.**
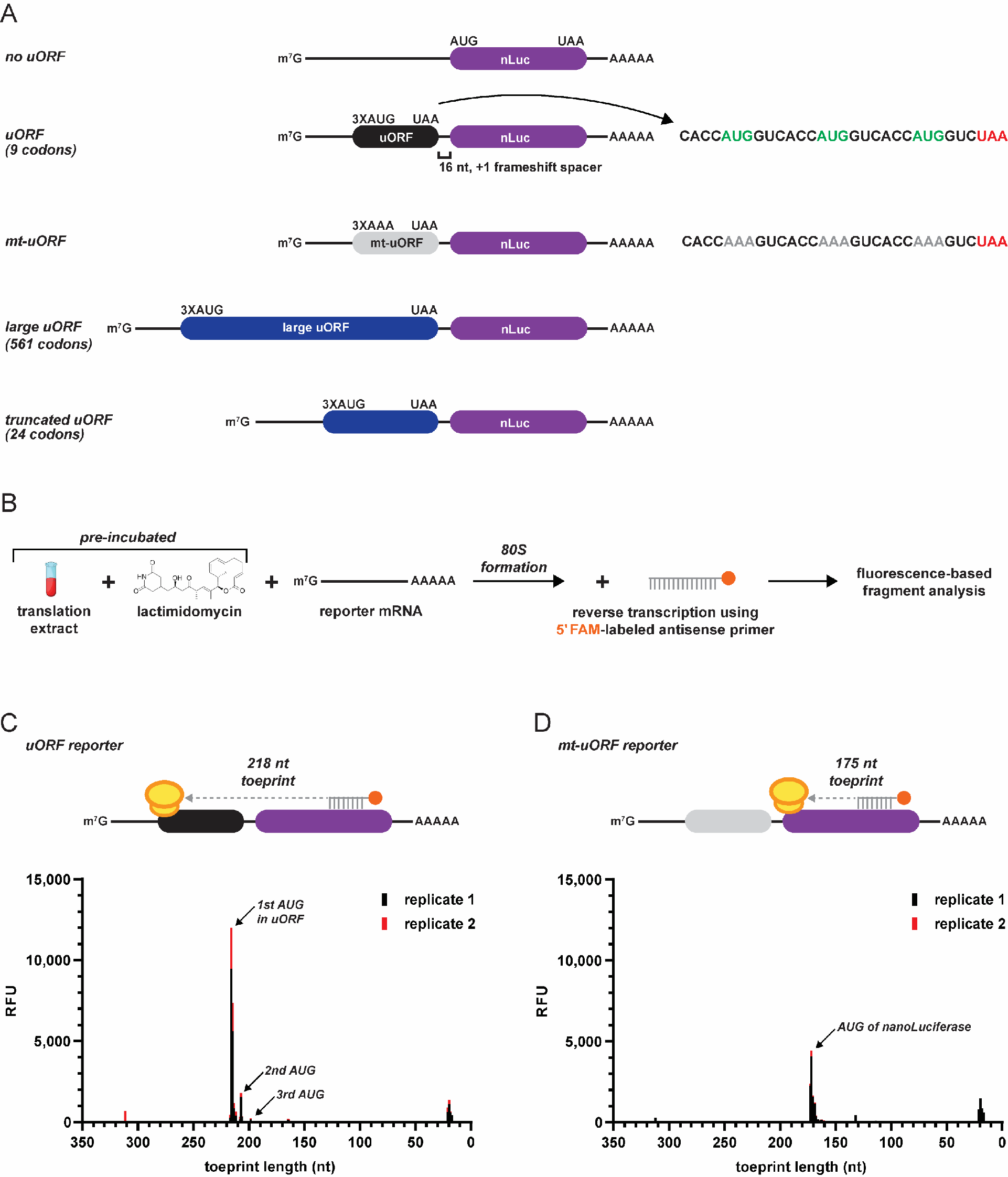
A small uORF with three start codons in perfect context is able to sequester all scanning initiation complexes *in vitro*. A) Design of re-initiation specific nanoLuciferase (nLuc) reporters used in this study. A 16 nt spacer between the variable-sized uORF and nLuc ORF allows specific detection of re-initiation. B) Schematic of ribosome toeprinting with FAM-labeled primers to detect sites of initiation with lactimidomycin pre-incubation. C-D) Ribosome toeprinting of 80S ribosomes after start codon recognition on (C) small uORF nLuc reporter mRNA and (D) mutated uORF nLuc reporter mRNA from *in vitro* translation. Signal from unused primer is seen at 20 nt. Signal from duplicate samples is shown in black and red.

We next used ribosome toeprinting to confirm that the 3XAUG uORF prevents leaky scanning and sequesters all detectable scanning PICs from the nLuc ORF (**Fig. 1B, C**). *In vitro* translation extracts were pre-incubated with lactimidomycin (binds to the E site of the 60S subunit) to inhibit the first translocation cycle of the 80S ribosome after initiation (Schneider-Poetsch et al. 2010). The 20 nt FAM-labeled reverse primer targeted a region in the nLuc coding sequence that was downstream enough to detect toeprints of 80S ribosomes at the AUG start codon of nLuc (175 nt), which would be present if leaky scanning past the uORF occurred, and at the AUG start codon of the uORF (218 nt). Indeed, for the uORF reporter mRNA, toeprint signal mapped primarily to the first and second AUG start codon of the uORF (218 and 210 nt, respectively), with some detectable signal mapping to the third AUG start codon (**Fig. 1C**).

Signal from unused primer is seen at 20 nt. Importantly, no signal was mapped to the start codon of nLuc (175 nt), which is in alignment with the design for the uORF to trap all scanning PICs.

As an additional control, we mutated the AUG start codons of the uORF to AAA (which does not support initiation) to allow all scanning PICs to bypass the uORF and initiate at the AUG start codon of nLuc (**Fig. 1A**). As expected, ribosome toeprints of this mutant uORF reporter produced signal that only mapped to the AUG start codon of nLuc (**Fig. 1D**). Similar results were seen with the uORF consisting of 10 consecutive AUG codons, but, because the start codons were not in optimal context, more initiation within the uORF was observed (**Supplemental Fig. 1A-C**). Lastly, by adapting an overlapping uORF design, our data demonstrates that the 3XAUG uORF captures 98-99% of scanning PICs in HeLa cells (**Supplemental Fig. 1D**). Together, these data support that the designed uORF essentially captures all detectable scanning PICs and allows all luciferase signal to be generated from re-initiation.

Consistent with uORFs generally being translational repressive elements, the small uORF repressed translation of nLuc 5-10-fold *in vitro* and in HeLa cells (**Fig. 2**). Translation of nLuc was rescued when the AUG start codons in the uORF were mutated to AAA codons.

**Figure 2.**
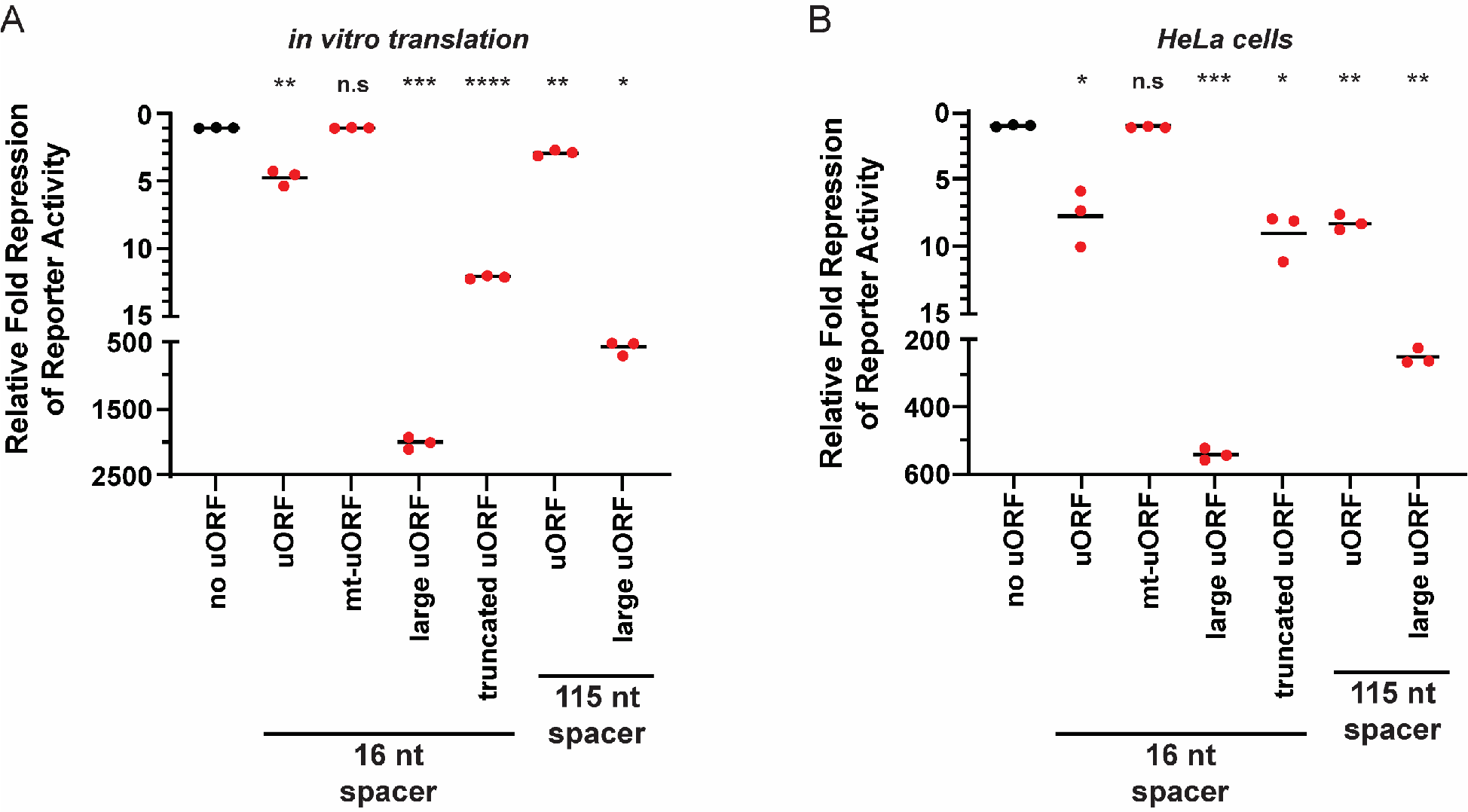
Translation re-initiation is more efficient after small uORFs *in vitro* and in HeLa cells. A-B) Response of nLuc reporters that harbor a small, mutant, large, or truncated uORF with a 16 nt or 115 nt intercistronic spacer from *in vitro* translation (A) and in HeLa cells (B). n=3 biological replicates. Bar represents the mean. Luminescence signals were set relative to the control no uORF reporter (in black). Comparisons against the control (in black) and experimental reporters (in red) were made using a two-tailed unpaired *t*-test with Welch’s correction. * = p<0.05. ** = p<0.05. *** = p<0.005. **** = p<0.0001. Exact p-values can be found in **Supplemental Table 2**.

Given that the average uORF in mammalian transcripts is much smaller than the main annotated protein coding ORFs (16 codons vs 460 codons) (Calvo et al. 2009), we tested how expanding the uORF length would affect re-initiation. The uORF was expanded from 9 codons to 561 codons by inserting the HaloTag-GFP (HT-GFP) coding sequence immediately downstream from the 3XAUG start codons and upstream of the stop codon. In both *in vitro* and in HeLa cells, the large uORF repressed translation of nLuc orders of magnitude more than the small uORF (**Fig. 2**). To confirm this observation was not due to differences in the sequence that the ribosome occupies during initiation and termination in the large uORF, we made a truncated HT-GFP uORF (total of 24 codons) that preserved the first and last 24 nt of the HT-GFP coding sequence (**Fig. 1A**). The truncated uORF rescued expression compared to the larger uORF *in vitro* and in HeLa cells (**Fig. 2A**). However, because the truncated uORF is still ∼3X larger than the small uORF, it was more repressive than the small uORF. Additionally, we tested whether increasing the spacer between the two cistrons encouraged re-initiation after small and large uORFs. Previous literature postulates that longer spacers (e.g., 115 nt vs. 16 nt use here) would allow more time for the scanning unrecycled small subunits to acquire eIFs and favor re-initiation (Kozak 1987; Vattem and Wek 2004). We found that a long 115 nt spacer between the cistrons decreased the repressive effect of only the large uORF in both *in vitro* and in HeLa cells (**Fig. 2**). The longer spacer was slightly effective *in vitro* with the 3XAUG uORF, but this minor effect was absent in HeLa cells (**Fig. 2**). It should be noted that re-initiation is generally an inefficient process; global analysis of termination revealed very few ribosomes are found downstream of stop codons (Wangen and Green 2020). Additionally, as seen in the prototypical uORF-containing mRNA, *GCN4*, a steady decline in scanning 43S PICs and elongating 80S ribosomes are seen across the 5ʹ UTR and among the four small uORFs, in both yeast and human cells (Wagner et al. 2020).

It remains possible that the greater repression of the large uORF could be due to the HT-GFP coding sequence forming an unexpected secondary structure that inhibited the ribosome or sequestered the 5ʹ cap. This seems unlikely because we obtained equivalent results with a different but equally large uORF sequence (**Supplemental Fig. 2**). Additionally, the sequence that could influence an initiating ribosome based off the known ribosome footprint size (∼30 nt) was preserved between the large and truncated uORFs. Nevertheless, we tested if the small and large uORF sequences produced equal levels of nLuc when fused to a P2A “ribosome skipping motif” and the nLuc ORF (**Supplemental Fig. 3**). The P2A motif allows the ribosome to release the nascent polypeptide but continue elongation (Szymczak and Vignali 2005; Liu et al. 2017). Thus, the same nLuc polypeptide from both fusion reporters is assayed. Signal from the large uORF-P2A-nLuc reporter was ∼4-fold lower than the control uORF-P2A- nLuc reporter *in vitro* (**Supplemental Fig. 3**). Importantly, the observed ∼4-fold difference does not rationally explain for the ∼400-fold difference in re-initiation between the small and large uORFs *in vitro* (**Fig. 2**). Together, these data support that large uORFs are more repressive than small uORFs because they allow less translation re-initiation.

**Figure 3.**
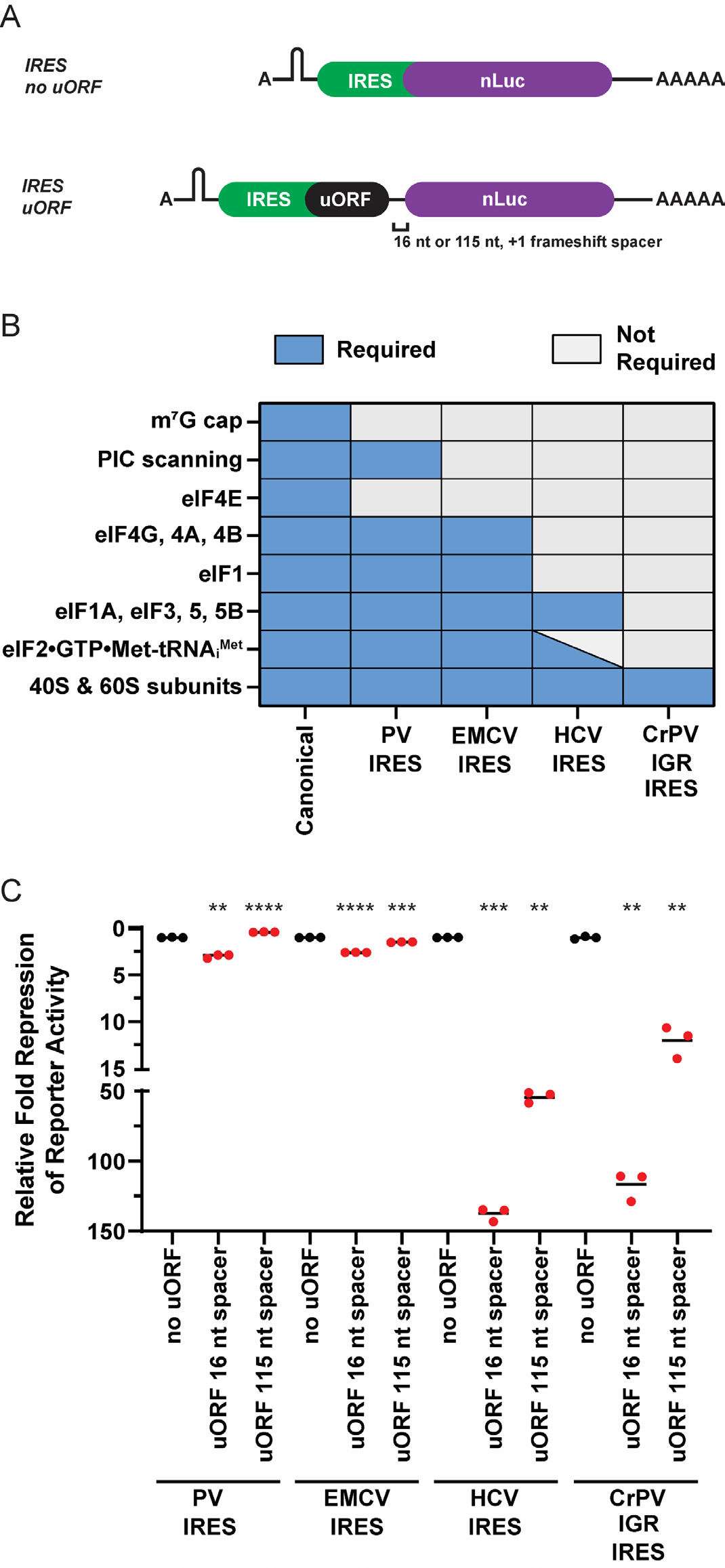
uORFs translated by IRESs that require less initiation factors permit less re- initiation. A) Schematic of A-capped IRES-mediated re-initiation reporters. A stable hairpin was inserted upstream of the IRES to block scanning ribosomes. B) Requirements of canonical initiation and class I-IV viral IRES-mediated initiation. C) Response of canonical initiation and viral IRES-dependent nLuc reporters without and with small uORFs from *in vitro* translation. n=3 biological replicates. Bar represents the mean. PV=poliovirus; EMCV=encephalomyocarditis virus; HCV=hepatitis C virus; CrPV IGR=cricket paralysis virus intergenic region; PIC=pre- initiation complex; eIF=eukaryotic initiation factor. Luminescence signal of each IRES uORF reporter (in red) was set relative to their respective control IRES no uORF reporter (in black). Comparisons between each control IRES and their respective experimental IRES reporters were made using a two-tailed unpaired *t*-test with Welch’s correction. * = p<0.05. ** = p<0.05. *** = p<0.005. **** = p<0.0001. Exact p-values can be found in **Supplemental Table 2**.

Recent reports using crosslinking and immunocapture of initiation factors (eIFs) have provided evidence that some eIFs may linger on the ribosome after initiation and could aid re-initiation if present after termination of a small uORF (Bohlen et al. 2020; Wagner et al. 2020). With this in mind, we next tested if re-initiation after small uORFs is as efficient if they are translated by ribosomes requiring less eIFs during initiation. We achieved this by taking advantage of class I-IV cap-independent viral internal ribosome entry sites (IRESs) that require subsets of eIFs (Pestova and Hellen 2003; Jackson et al. 2010; Jaafar et al. 2016; Petrov et al. 2016) (**Fig. 3A, B**). In certain cases, IRES can stimulate initiation without requiring any eIFs or the initiator tRNA (**Fig. 3A, B**). We consistently found that IRES-mediated translation is less efficient than canonical translation and we were only able to test the effect of the small uORF and still have luciferase signal above background in the linear range (data not shown). In alignment with the model that some eIFs can stay bound to elongating 80S ribosomes and aid in re-initiation (Bohlen et al. 2020; Wagner et al. 2020), we observed that the small uORFs repressed translation greater when the IRES utilized less initiation factors (**Fig. 3B, C**). We next asked how increasing the spacer between the IRES uORF and nLuc from 16 nt to 115 nt affected translational repression and re-initiation. If the longer spacer permitted the scanning unrecycled small subunit more time to acquire eIFs, we would expect that the IRESs that utilize less eIFs to be the most affected. Indeed, increasing the spacer blunted the repressive nature of uORFs when translated via the Hepatitis C Virus (HCV) IRES and Cricket Paralysis Virus Intergenic Region (CrPV IGR) IRES (**Fig. 3C**).

The presence of uORFs presents a major challenge to mRNAs as termination in the 5ʹ UTR resembles how ribosomes recognize deleterious premature termination codons (PTCs) through nonsense-mediated mRNA decay (NMD). Termination ≥50 nts upstream of an exon junction complex (EJC) typically robustly triggers NMD in mammalian cells (Lykke-Andersen and Jensen 2015; Yi et al. 2021). However, termination at uORFs almost certainly occurs upstream of an EJC if the uORF is translated during the pioneering round of translation. Others have postulated that re-initiation could be a method that ribosomes use to bypass triggering NMD at PTCs within the major ORF (Jagannathan and Bradley 2016; Dyle et al. 2020). We next asked if an mRNA with a translated small uORF that greatly favors re-initiation is as proportionally stable as an mRNA with a translated large uORF that drastically disfavors re-initiation (**Fig. 2 and Supplemental Fig. 2**). To answer this question, we used a Tet-Off system to selectively turn off reporter transcription and measured reporter mRNA levels over an 8 hr time course after the addition of doxycycline. In this experiment, we used the same reporter design as described in **Fig. 1**, to capture essentially all scanning 43S PICs in HeLa cells (**Supplemental Fig. 1D**), but with the addition of a small functional chimeric intron in the nLuc ORF. As a positive control for enhanced mRNA decay, we included a reporter that harbors an intron in the 3ʹ UTR greater than 50 nt downstream from the stop codon of nLuc, which should stimulate NMD and have a shorter mRNA half-life. As expected for stimulating NMD, the no uORF + 3ʹ UTR intron reporter had a ∼2hr shorter half-life than the control no uORF reporter (3.71 ± 0.52 hr vs. 5.84 ± 1.12 hr, respectively) (**Fig. 4**). The small and large uORF reporters had even shorter mRNA half-lives of 3.16 ± 0.34 hr and 2.49 ± 0.33 hr, respectively (**Fig. 4**).

**Figure 4.**
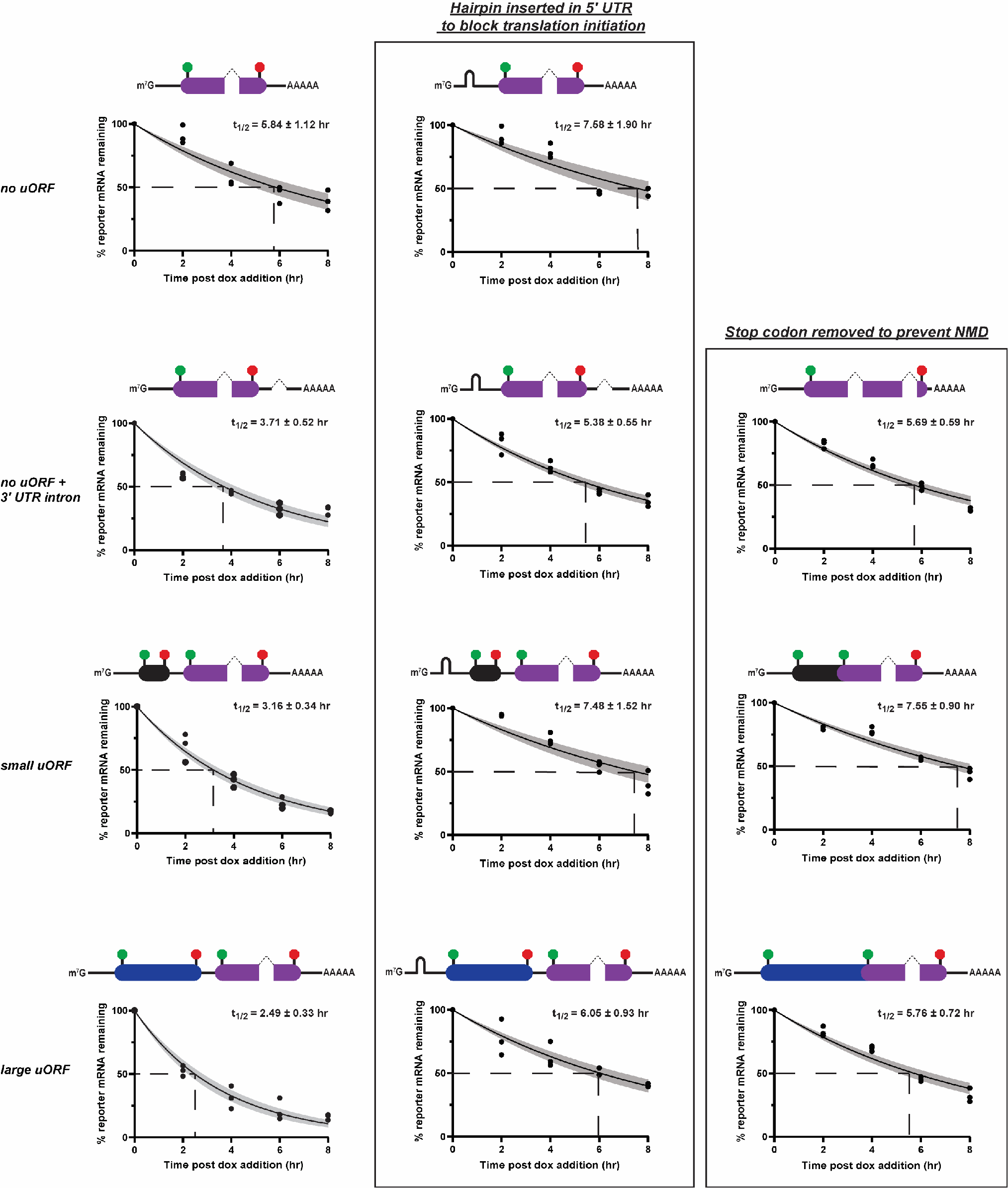
Small and large uORFs both robustly stimulate mRNA decay in HeLa cells. A Tet-Off system triggered with 2 µg/mL doxycycline (dox) was used to determine reporter mRNA half-lives in HeLa cells. Schematic of reporter is located above each mRNA decay curve. The small and large uORFs (same design as in Fig. 1) are shown in black and blue, respectively.nLuc ORF is in purple. Start and stop codons are green and red circles, respectively. Introns in the nLuc ORF and 3ʹ UTR are shown as dotted angled lines. As controls to rescue mRNA half-life from NMD, a strong hairpin in the 5ʹ UTR was inserted to limit translation initiation (middle column) and stop codons were mutated from UAA to UAC to prevent termination upstream of a spliced intron (right column). n=3 biological replicates. A non-linear regression was used to calculate the mRNA half-lives and is shown as the line with the 95% confidence interval included as a watermark. The half-life is reported with the error for the 95% confidence interval range.

Despite harboring drastically different re-initiation capabilities (**Fig. 2B**), the half-life of the small uORF reporter was only ∼40 min longer than the large uORF reporter but still substantially shorter than the no uORF control (**Fig. 4**). These data suggest that although the small uORF promotes re-initiation ∼100-fold better than the large uORF in cells (**Fig. 2B**), this high level of re-initiation after the small uORF does not robustly allow the mRNA to evade NMD, and that re-initiation after uORFs minorly contributes to stability. Importantly and as expected for NMD substrates, the shortened half-life of the small and large uORF reporters compared to the control no uORF reporter was completely rescued when translation initiation was limited by the presence of a strong hairpin in the 5ʹ UTR and when translation termination in the 5ʹ UTR was abolished by mutation of the stop codon of the uORF from UAA to UAC (**Fig. 4**). Rescue of mRNA stability for the no uORF + 3ʹ UTR intron reporter (NMD positive substrate control) was also observed when the same targeted approaches to limit NMD were implemented (**Fig. 4**).

Finally, we further investigated the connection between uORF length and mRNA stability on a transcriptome-wide scale by mining published BRIC-seq datasets (Tani et al. 2012; Maekawa et al. 2015) and determined the cumulative predicted uORF length for each mRNA. We found no statistical difference between the mean mRNA half-life of transcripts that contained varying cumulative computationally predicted uORF lengths in multiple data sets (**Supplemental Fig. 4**). However, transcripts without an uORF were on average more stable and had longer half-lives (**Supplemental Fig. 4**). Collectively, these data suggest translation re-initiation after uORFs is not a robust method for mRNAs to prevent NMD, and that re-initiation after uORFs provides less stabilization than previously thought. These data are consistent with the general mammalian NMD model that termination upstream of an EJC triggers NMD and that re-initiation by definition can only occur after termination is completed. However, ribosome readthrough, which occurs when a near-cognate aminoacyl tRNA successfully competes against eRF1•eRF3•GTP for the A site at stop codons, prevents termination and can noticeably stabilize NMD targets (Annibaldis et al. 2020; Embree et al. 2022). Readthrough is rather inefficient but can be made more competent by aminoglycosides and other small molecules (Martin et al. 2014; Wangen and Green 2020; Embree et al. 2022).

In total, our data supports a model where, in mammalian cells, uORF length highly controls re-initiation but only modestly affects termination-dependent decay in the 5ʹ UTR, and that the decision of whether NMD is stimulated occurs before re-initiation. Whether a ribosome actively translates an uORF and successfully terminates upstream of an EJC is the primary influence if uORFs stimulate mRNA decay. The coding capacity of a single mRNA can be increased with larger uORFs, but a trade-off ensues as they do not favor re-initiation and provide less translational regulation. This control can be regulated by leaky scanning, particularly if the uORF harbors a start codon in sub-optimal Kozak context. For example, many stress response mRNAs harbor uORFs and are targeted by NMD during normal growth conditions; upon cell stress, the uORFs are bypassed and are not translated, resulting in the abundance of stress response mRNAs to increase (Gardner 2008; Karam et al. 2015; Young and Wek 2016; Wek 2018). Leaky scanning past uORFs would allow elongating ribosomes on the main ORF to displace EJCs and Upf1, effectively preventing subsequent uORF translation from stimulating NMD. Clearly, mammalian evolution has not favored large uORFs as the average mammalian uORF is 16 codons (Calvo et al. 2009). This may be partially explained not only by the fact that large uORFs provide more opportunity for other translation-dependent decay pathways (e.g., codon-optimality-mediated decay, no-go decay from ribosome collisions), but they also prevent translation of the downstream main ORF by decreased re-initiation.

## MATERIALS AND METHODS

### Plasmids

Complete sequences of reporter plasmid inserts are located in the Supplementary Material. All plasmids were derived from previously described pcDNA3.1(+)/AUG-nLuc-3XFLAG and pcDNA3.1-D/CrPV IGR IRES nLuc-3XFLAG (Kearse et al. 2016). The CrPV IGR IRES reporter was additionally modified to contain a strong hairpin upstream of IRES element to block scanning pre-initiation complexes. The HT-GFP ORF was taken from pHaloTag-EGFP (a gift from Thomas Leonard and Ivan Yudushkin; Addgene plasmid # 86629). pGL4.13 (encodes Firefly Luciferase [FFLuc]) was obtained from Promega (# E6681). IRES-containing nLuc reporters were generated using an overlapping PCR method and cloned into pcDNA3.1(+) or pcDNA3-1D. The PV IRES template was pcDNA3 RLUC POLIRES FLUC and was a gift from Nahum Sonenberg (Addgene plasmid # 45642). The EMCV IRES and HCV IRES templates were kind gifts from Aaron Goldstrohm. All IRES reporters contained the same strong hairpin upstream of the IRES element. 5ʹ UTRs, uORFs, introns, hairpins, and mutations were introduced using the Q5 Site-Directed Mutagenesis Kit (NEB # E0554S) or were synthesized by Genscript. Complete sequences for all reporters can be found in the Supplemental Material.

To make pTet-Off All-In-One plasmids, pcDNA3.1(+)/no uORF nLuc plasmid was subjected to two rounds of mutagenesis using the NEBuilder HiFi DNA Assembly Master Mix with 25 bp overhangs. First, the complete CMV promoter was replaced with the tetracycline- responsive PTight promoter from pCW57.1-MAT2A (a gift from David Sabatini; Addgene plasmid # 100521). Second, the neomycin resistance gene coding sequence was replaced with the tTA- Advanced coding sequence from pCW57.1-MAT2A. The different uORF nLuc inserts were then subcloned into this pTet-Off All-In-One backbone at SacI and XbaI sites. The 133 bp chimeric intron from pCI-neo (Promega # E1841) was inserted into the nLuc ORF by using the Q5 Site-Directed Mutagenesis Kit.

All oligonucleotides were obtained from Integrated DNA Technologies. TOP10 *E. coli* cells were used for all plasmid propagation and cloning. Reporters and any mutated sites were fully Sanger sequenced at The Ohio State Comprehensive Cancer Center Genomics Shared Resource (OSUCCC GSR).

### *In vitro* transcription

Reporter mRNAs were synthesized using linearized plasmids as templates for run off transcription with T7 RNA polymerase as previously described (Kearse et al. 2016) with the single exception that XbaI was used to linearize all plasmids. All mRNAs were transcribed at 30°C for 2 hrs using the HiScribe T7 High Yield RNA Synthesis Kit (NEB # E2040S) and were co-transcriptionally capped (8:1 cap analog to GTP for ∼90% capping efficiency) and post-transcriptionally polyadenylated. Non-IRES mRNAs were capped the 3’-O-Me-m7G(5’)ppp(5’)G anti-reverse cap analog (NEB # S1411L). IRES mRNAs were capped with the A(5’)ppp(5’)G cap analog (NEB # S1406L). Post-transcriptional polyadenylation was performed using *E. coli* Poly(A) Polymerase (NEB # M0276L). mRNAs were purified using the Zymo RNA Clean and Concentrator-25 (Zymo Research # R1018), eluted in RNase-free water, aliquoted in single use volumes and stored at -80°C.

### *In vitro* translation and luciferase assay

10 μL *in vitro* translation reactions were performed in the linear range using 3 nM mRNA in the Flexi Rabbit Reticulocyte Lysate (RRL) System (Promega # L4540) with final concentrations of reagents at 30% RRL, 10 µM amino acid mix minus Leucine, 10 µM amino acid mix minus Methionine, 0.5 mM MgOAc, 100 mM KCl, and the addition of 8 U murine RNase inhibitor (NEB # M0314L) (Kearse et al. 2016). Reactions were incubated for 30 min at 30°C, terminated by incubation on ice, and then diluted with 40 µL Glo Lysis Buffer (Promega # E2661). 25 µL of diluted reaction was mixed with 25 µL of prepared Nano-Glo Luciferase Assay System (Promega # N1120) for 5 min in the dark on an orbital shaker. Luminescence was measured using a Promega GloMax Discover Microplate Reader. Reactions testing IRES nLuc reporters were additionally supplemented with 50 ng/µL (final) competitor FFLuc mRNA (TriLink Biotechnologies # L-7602-100) to increase the fidelity of the IRES nLuc signal; addition of competitor mRNA did not hinder re-initiation levels of the canonically translated 3XAUG uORF nLuc mRNA (data not shown). We have previously shown that RRL translation reactions with 3 nM mRNA input (final) incubated at 30°C for 30 min is in the dynamic linear range for a wide variety of reporters (Kearse et al. 2016).

### Fluorescent ribosome toeprinting

60 µL *in vitro* translation RRL reactions (same final concentrations of reagents as above except for mRNA) were pre-incubated with 50 µM lactimidomycin (Sigma # 5062910001; 5 mM stock in DMSO) for 10 min at 30°C then placed on ice. 25 nM capped and polyadenylated mRNA was added (increased concentration was important to detect weaker signals), gently mixed, and incubated for an additional 10 min at 30°C to allow inhibited 80S ribosomes to form after start codon recognition. To each reaction, 20 µL 5X AMV RT buffer (final: 50mM Tris-HCl (pH 8.3), 50 mM KCl, 10 mM MgCl2, 0.5 mM spermidine, 10 mM DTT), 10 µL 10 µM 5’-FAM labeled-reverse primer (20 nt), 2 µL 25mM complete dNTP set (final of each dNTP at 0.5 mM), 6 µL nuclease-free water, and 2 µL AMV Reverse Transcriptase (stock at 20-25 U/µL) was added and reverse transcriptase (RT) was allowed to progress for 35 min at 30°C. The higher MgCl2 concentration in the RT reaction inhibits new initiation complex formation. Control reactions with 25 mM reporter mRNA in water were treated identically and were used to determine background from the RT reaction. FAM-labeled cDNA was extracted by transferring the 100 µL RT reaction to a new microcentrifuge tube with 150 µL nuclease-free water and adding 250 µL saturated Phenol:Chloroform:Isoamyl Alcohol (25:24:1), pH 8. After vigorous mixing for 1 min, samples were centrifuged at room temperature at 16,000 rcf for 5 min. The top aqueous phase was transferred, and re-extracted with saturated Phenol:Chloroform:Isoamyl Alcohol (25:24:1), pH 8. The final aqueous supernatant was then concentrated using the Zymo DNA Clean and Concentrator-5 using a 7:1 ratio following the manufacture’s recommendation. FAM-labeled cDNA was eluted in 7 µL nuclease-free water. 5 µL of each eluate was mixed with 10 µL Hi-Di Formamide (Thermo # 4440753), spiked with a LIZ 500 size standard, and subjected to fragment analysis using Applied Biosystems 3130xl Genetic Analyzer with POP-7 polymer with all fragments being reported. To determine which signals were caused by inhibited 80S ribosomes at start codons, signal from the control samples (RNA in water + RT reaction) were subtracted from the reactions with RRL and inhibitor. Primer sequence is included in the **Supplemental Table 1**.

### Cell culture, transfection, and luciferase assay

HeLa cells were obtained from ATCC and maintained in high glucose DMEM supplemented with 10% heat-inactivated FBS, 1% penicillin-streptomycin, and 1% non-essential amino acids in standard tissue culture-treated plastics. HeLa cells were seeded 24 hr before transfection so that on the day of transfection they were at 50% confluency. ViaFect (Promega # E4982) was used at a 3:1 ratio with 1 µg total plasmid (500 ng nLuc plasmid + 500 ng pGL4.13) in 100 µL in Opti-MEM (Thermo Fisher # 31985062). For 96-well plates, HeLa cells were transfected with a total of 100 ng (10 µL of the transfection mix). 24 hr post transfection, media was aspirated, and cells were lysed in 100 µL Glo Lysis Buffer (Promega # E2661) for 10 min on an orbital shaker. 25 µL of lysate was then mixed with 25 µL of ONE-Glo (Promega # E6120) or 25 µL of prepared Nano-Glo Luciferase Assay System (Promega # N1120) and detected as described above. nLuc signal was then normalized to FFLuc signal of the same sample to normalize for transfection efficiency. Statistical analysis results with exact p-values for the main figures can be found in **Supplemental Table 2**; all Supplemental Figures contain exact p-values.

### Tet-Off system and mRNA decay measurements

HeLa cells were seeded and maintained in complete media as described above, but supplemented with 10% Tet-approved FBS (Thermo Fisher # A47364-01). 24 hr post seeding in a 10 cm plate, 50% confluent cells were transfected with 6 µg of total plasmid (3 µg pTet-Off All-In-One plasmids + 3 µg pGL4.13) using ViaFect. 24 hr post transfection, cells were trypsinized, diluted, and seeded in five 12-well dishes. 48 hrs later, when cells were ∼75% confluent, media was replaced with media containing 2 µg/mL doxycycline (MP Biomedicals # 195044) (stock at 1 mg/mL in water) in the dark. At the indicated time points, total RNA was extracted using TRIzol (Thermo Fisher # 15596018) following the manufacture’s recommendations. 500 ng of total RNA was then DNase-treated with amplification grade DNase (Thermo Fisher # 18068015). The entire final 11 µL DNase reaction was then used to synthesize cDNA with oligo-dT primers and random hexamers using the iScript Reverse Transcription Supermix for RT-qPCR (Bio-Rad # 1708841) following the manufacture’s protocol. cDNA was then diluted 1:100 and 1.5 µL was use per 15 µL reaction with iTaq Universal SYBR Green Supermix (Bio-Rad # 1725122) and 250 nM primers (final) on a Bio-Rad CFX Connect Real-Time PCR Detection System using Bio-Rad CFX Maestro software to calculate expression levels. Reporter levels were normalized to RPS17 and half-lives were calculated using first order exponential decay trend lines, calculated by non-linear regression in GraphPad Prism 9.1.2. The error for the 95% confidence interval range were plotted along the mean of three biological replicates. Reverse transcriptase minus reactions were used to confirm less than 2% of reporter signal is from contaminating plasmid DNA. All primer sequences are available in **Supplemental Table 1**.

### Bioinformatic analysis of uORF length and mRNA decay

mRNA half-lives measured using BRIC-seq was obtained from published literature (Tani et al. 2012; Maekawa et al. 2015; Imamachi et al. 2017). Custom Python scripts were written to calculate predicted uORFs in each mRNA, add them together, and then assigned it to the previously defined mRNA half-life. If mRNAs had multiple uORFs, the lengths of all uORFs were summed together for a cumulative computationally predicted uORF length. Both scripts have been deposited in GibHub (github.com/michaelkearse/uORF_Half-life). The scripts utilize the RefSeq Transcript and RefSeq Reference Genome Annotation files for human genome build 38 from NCBI for reference. If a gene had multiple transcripts listed, then the longest transcript (usually isoform 1) was used. Only AUG-encoded uORFs were determined. Transcripts were then binned by cumulative uORF length in five codon increments with at least 100 mRNAs in each bin. The total sample number per bin is included in **Supplemental Table 3**, along with the cumulative computationally predicted uORF length and half-life for each mRNA.

## ACKNOWLEDGEMENTS

Experiments were conceived and performed by PJR and MGK with input and help from EPB. The bioinformatics analysis was performed by JAS with input from AHMB. The manuscript was written by PJR and MGK with input from JAS, EPB, and AHMB. PJR was supported by NIH grants T32GM086252 and T32GM141955. This work was supported by NIH grants R00GM126064 (to MGK) and R35GM146924 (to MGK). We thank members of the Kearse lab for critically reading the manuscript, as well as Amrit Singh, Kurt Fredrick, and Karin Musier-Forsyth for stimulating discussions. We also thank Christine Daugherty at The Ohio State Comprehensive Cancer Center Genomics Shared Resource (OSUCCC GSR) for her critical help with the fluorescent toeprinting analysis. The OSUCCC GSR is supported by NIH grant P30CA016058.

---

**Supplemental Figure 1.**
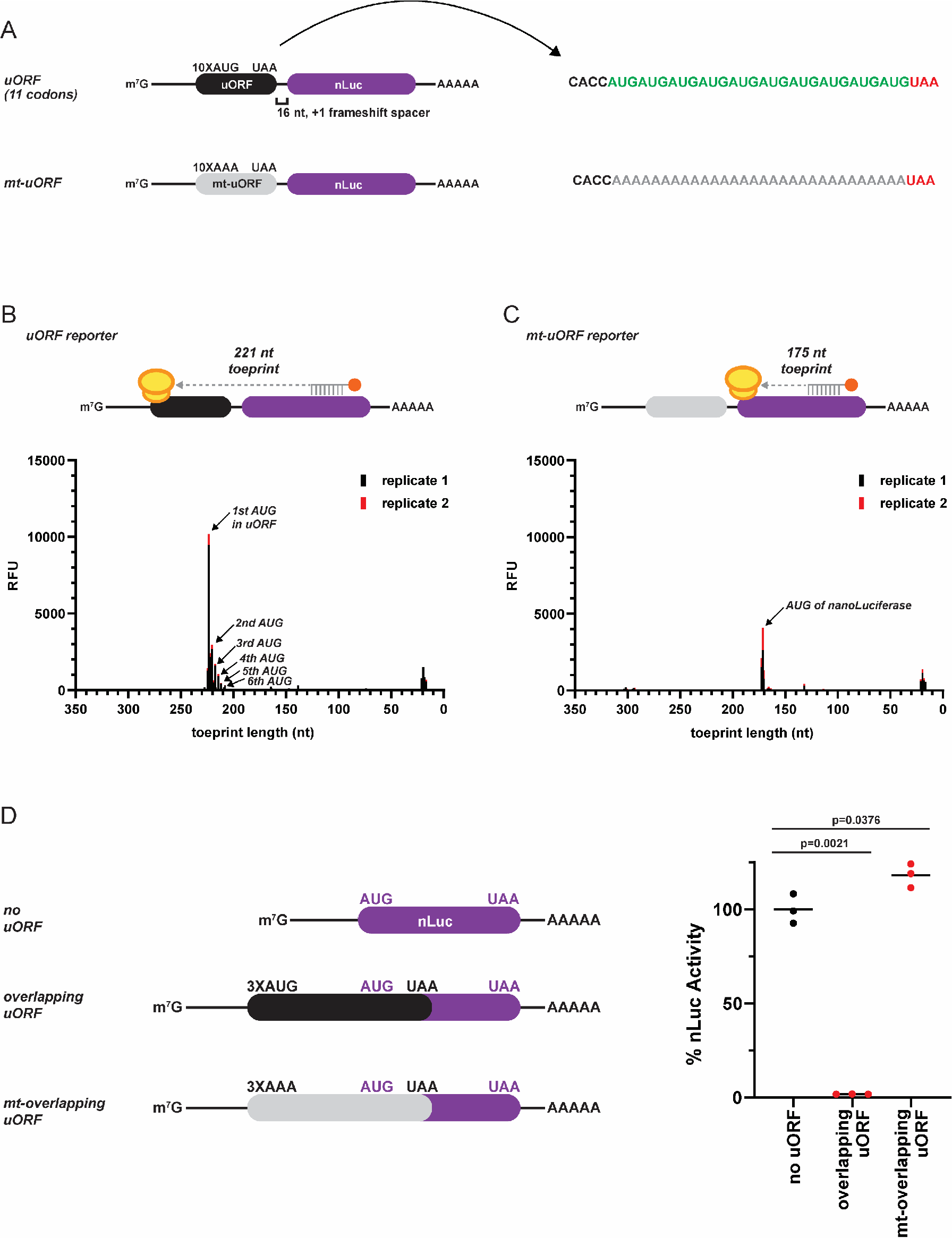
Confirming the efficacy of designed uORFs to sequester all scanning initiation complexes *in vitro* and in HeLa cells. A) Design of re-initiation specific nanoLuciferase (nLuc) reporters harboring an uORF encoded by 10 consecutive AUG start codons (uORF) or AAA codons (mt-uORF). A 16 nt spacer between the uORF and nLuc ORF allows specific detection of re-initiation. B-C) Ribosome toeprinting of lactimidomycin inhibited 80S ribosomes after start codon recognition on (B) small uORF nLuc reporter mRNA and (C) mutated uORF nLuc reporter mRNA from *in vitro* translation. Signal from duplicate samples is shown in black and red. D) Response of nLuc reporters that harbor an overlapping 3XAUG start codon uORF in HeLa cells. Here, the stop codon of the uORF (black) is downstream and out-of-frame of the start codon of nLuc (purple). Thus, active nLuc is only produced if a 43S PIC scans through the overlapping uORF (“leaky scanning”) and initiates at the start codon of nLuc. n=3 biological replicates. Bar represents the mean. Comparisons were made using a two-tailed unpaired *t*-test with Welch’s correction.

**Supplemental Figure 2.**
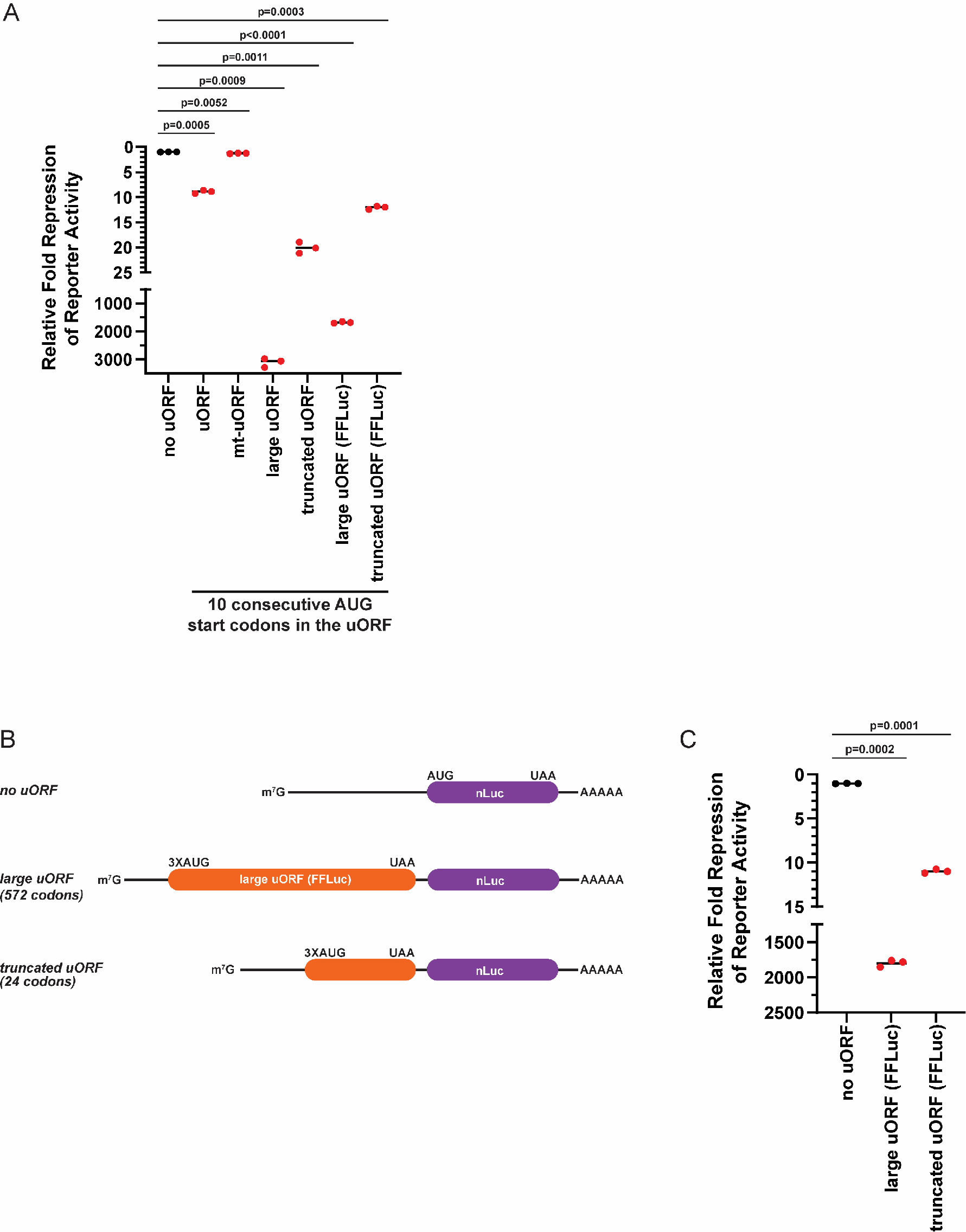
Less translation re-initiation after large uORFs is not strictly uORF sequence specific. A) Response of nLuc reporters that harbor a small, mutant, large, or truncated uORF from *in vitro* translation. All uORFs had 10 consecutive AUG start codons as described in **Supplemental Figure 1A** (instead of the three AUG start codons in perfect Kozak context in **Fig. 1**). Large and truncated uORFs using the FFLuc sequence are clearly labeled from those that used the original HT-GFP sequence. n=3 biological replicates. Bar represents the mean. Comparisons were made using a two-tailed unpaired *t*-test with Welch’s correction. B) The large uORF sequence described in **Fig. 1** was switched from HT-GFP to Firefly Luciferase (FFLuc). Three AUG start codons in perfect Kozak context were used to trap all scanning initiation complexes at the uORF. C) Response of nLuc reporters that harbor a large (FFLuc) or truncated uORF (FFLuc) from *in vitro* translation. Similar repression and rescue were seen here with FFLuc as the large uORF as observed with HT-GFP sequence in **Fig. 2**. n=3 biological replicates. Bar represents the mean. Comparisons were made using a two-tailed unpaired *t*-test with Welch’s correction.

**Supplemental Figure 3.**
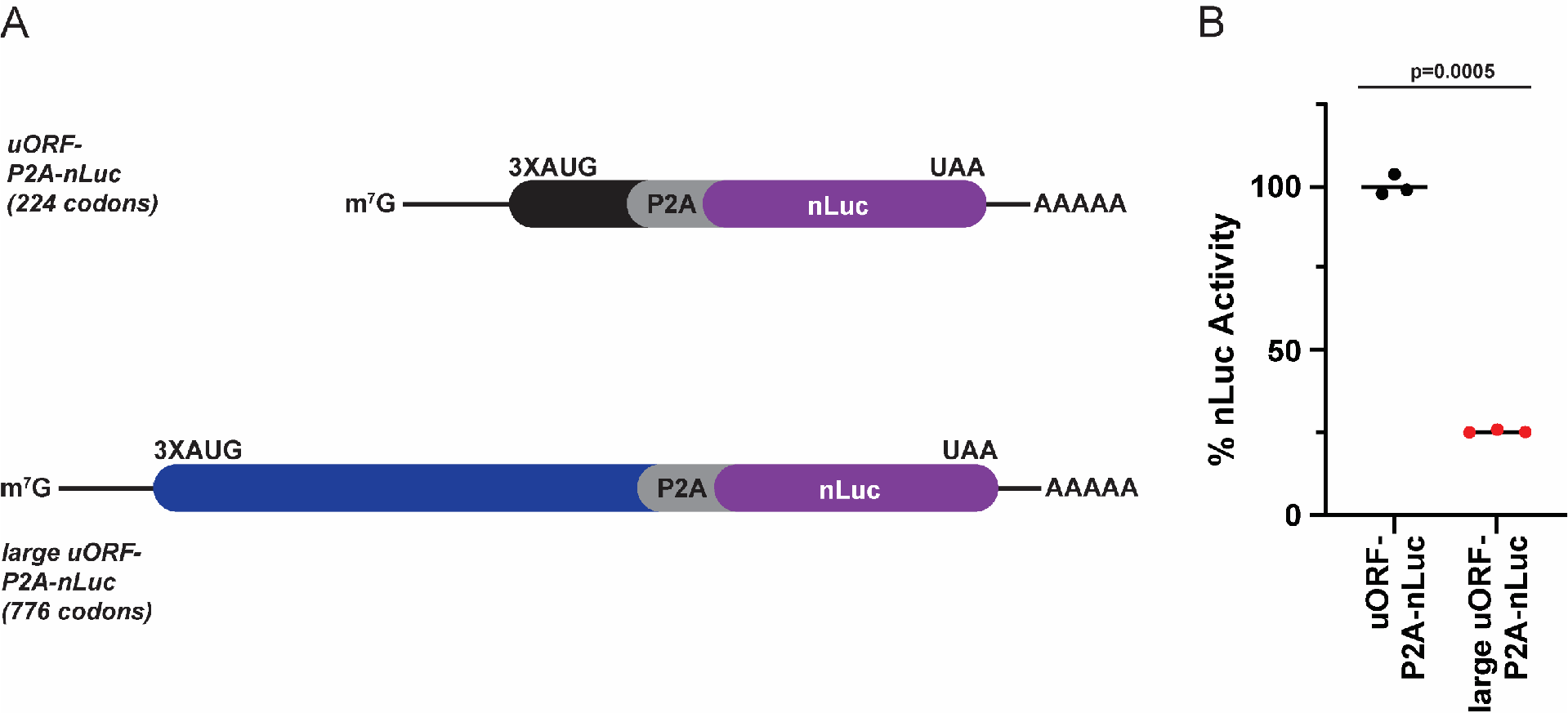
Small and large uORF P2A fusion reporters to test translation efficiency *in vitro*. A) Insertion of the P2A “ribosome skipping motif” (gray) was used to assess the relative translation efficiency of reporters that harbored a 3XAUG start codon sequence without and with the large HT-GFP (blue) sequence upstream of the nLuc coding sequence (purple). B) Relative nLuc activity of the small and larger uORF P2A fusion reporters from *in vitro* translation. n=3 biological replicates. Bar represents the mean. Comparisons were made using a two-tailed unpaired *t*-test with Welch’s correction.

**Supplemental Figure 4.**
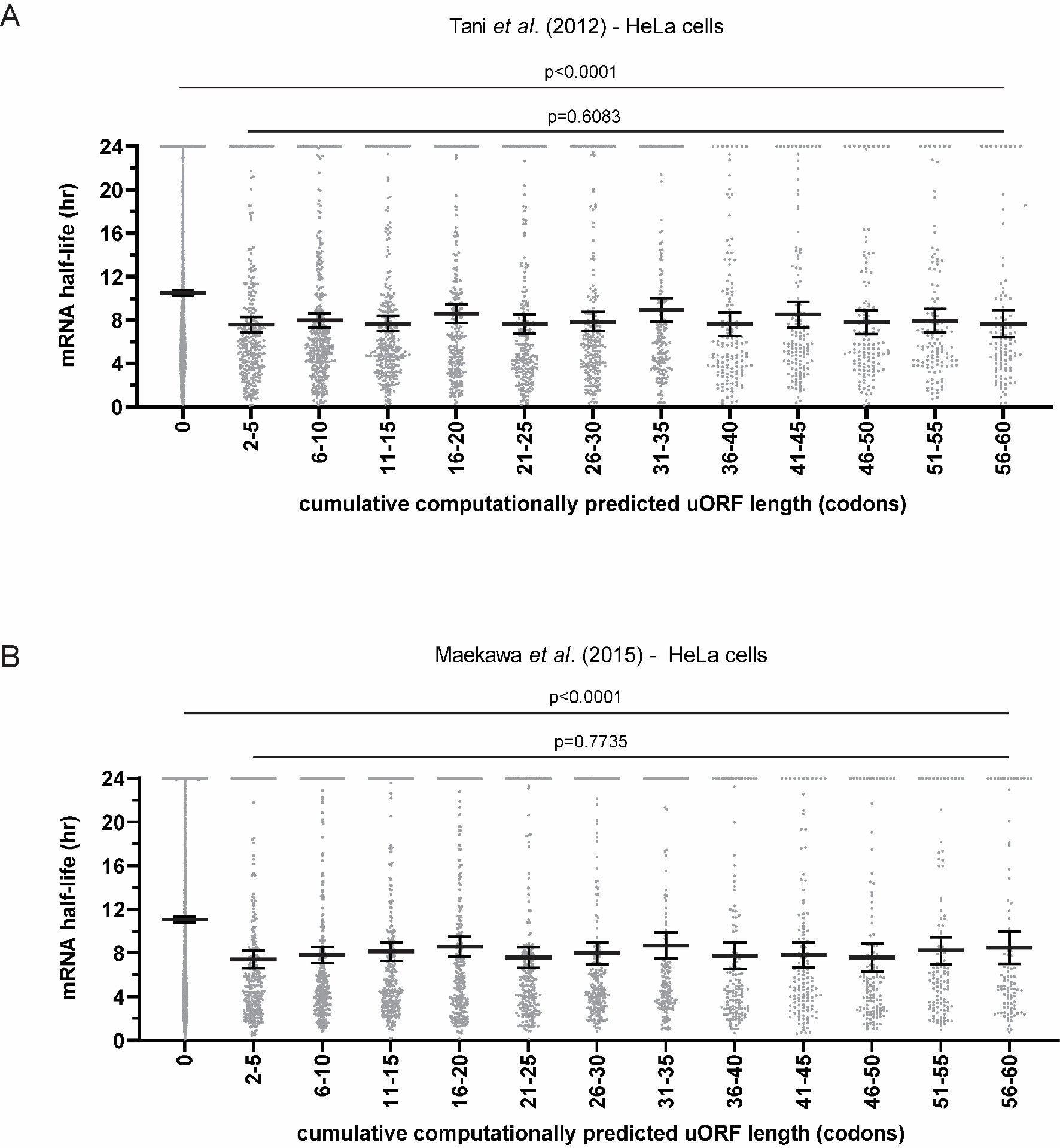
uORF length has minimal effect on mRNA stability in cells. A-B) Transcriptome-wide comparison of cumulative computationally predicted uORF length and mRNA half-life from Tani *et al*. (2012) (A) and Maekawa *et al*. (2015) (B). Bar represents the mean ± 95% confidence interval. One-way Welch’s ANOVA was used to compare between cumulative uORF length bins.

## SUPPLEMENTAL MATERIAL

Supplemental Table 1. Primers used in this study

Supplemental Table 2. List of p-values for each comparison made in Figures 2 and 3

Supplemental Table 3. List of cumulative uORF length and mRNA half-life re-analyzed from Tani et al. (2012) and Maekawa et al. (2015).

## Sequences of reporters used in this study

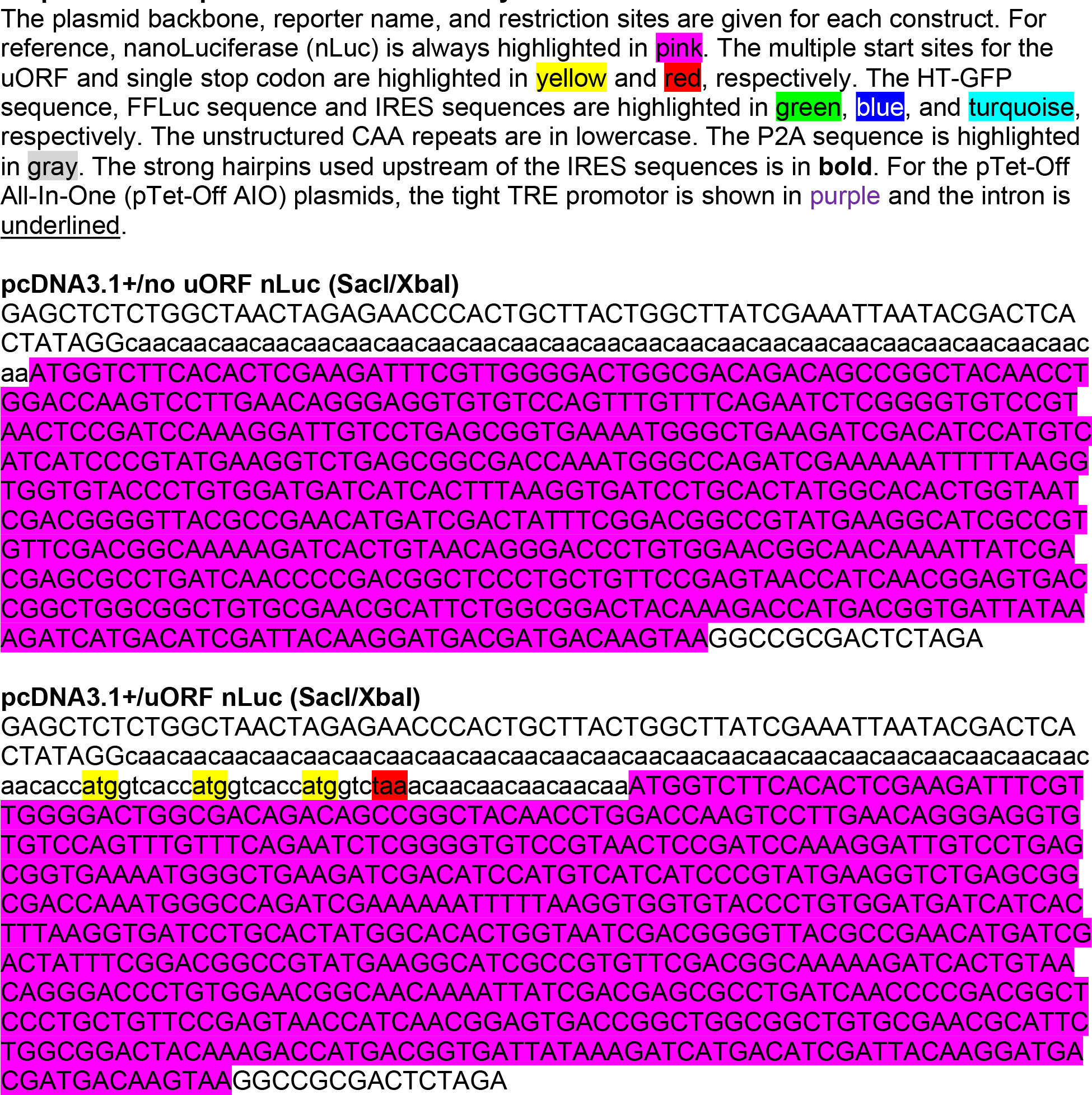
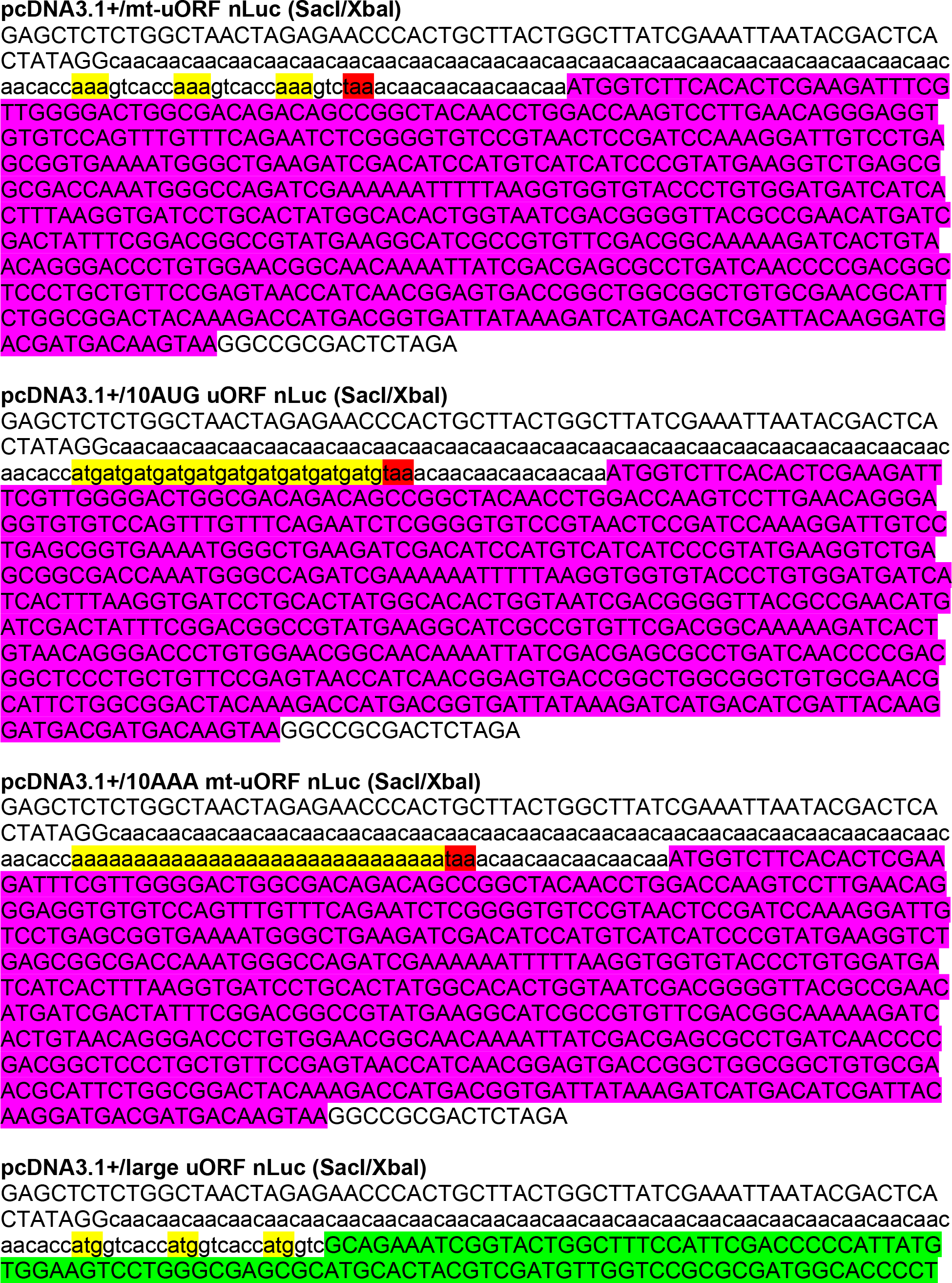
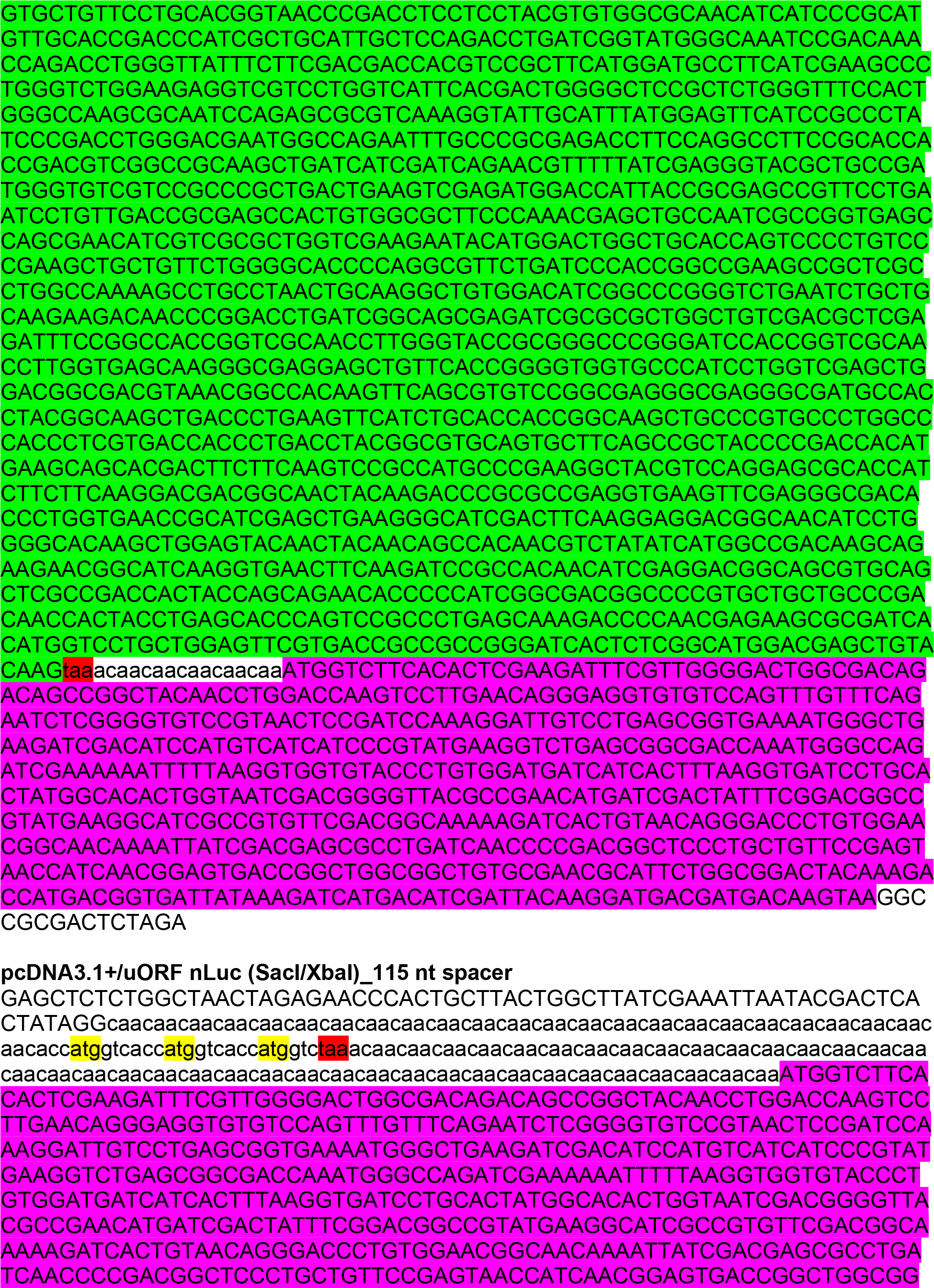
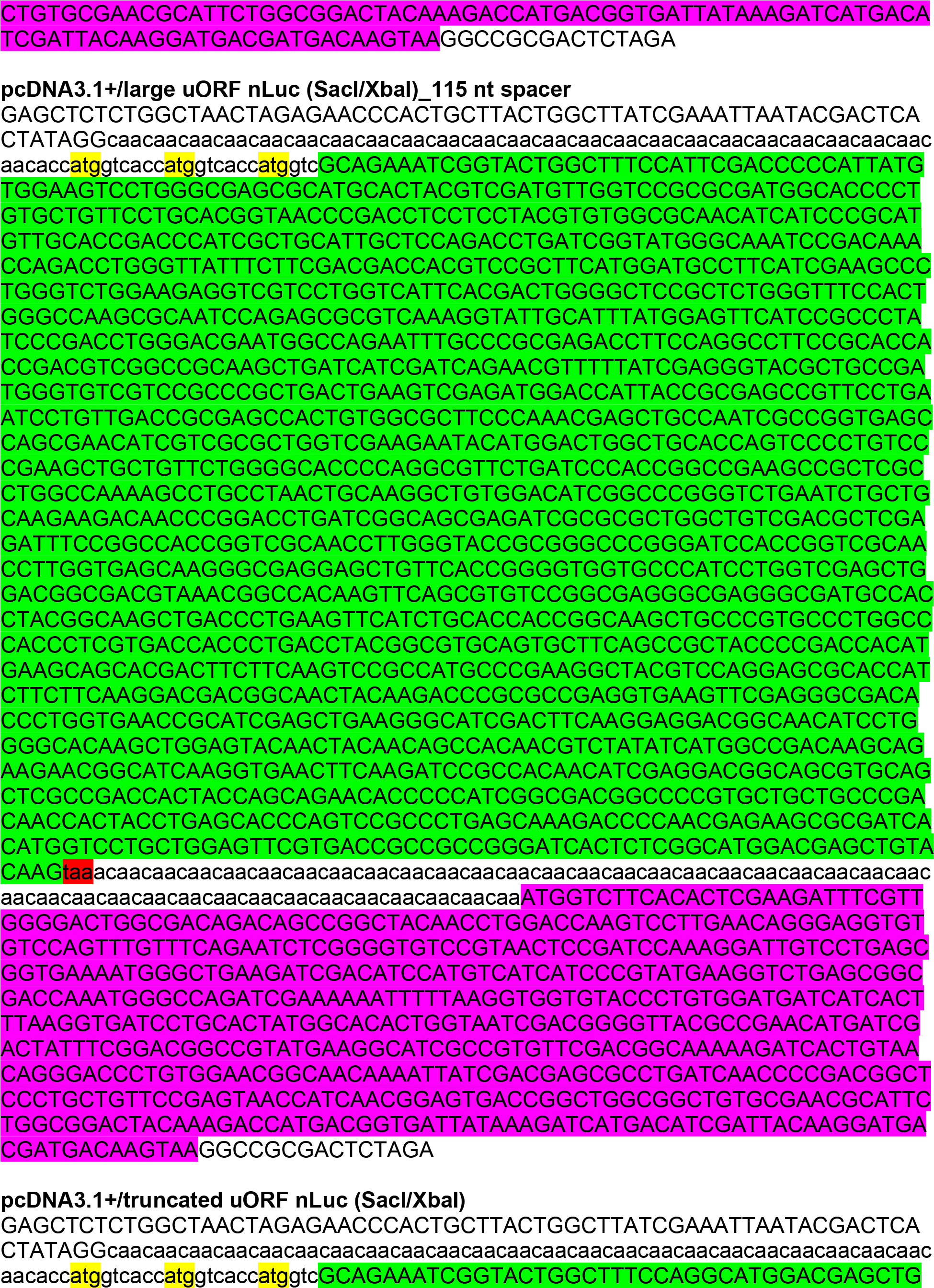
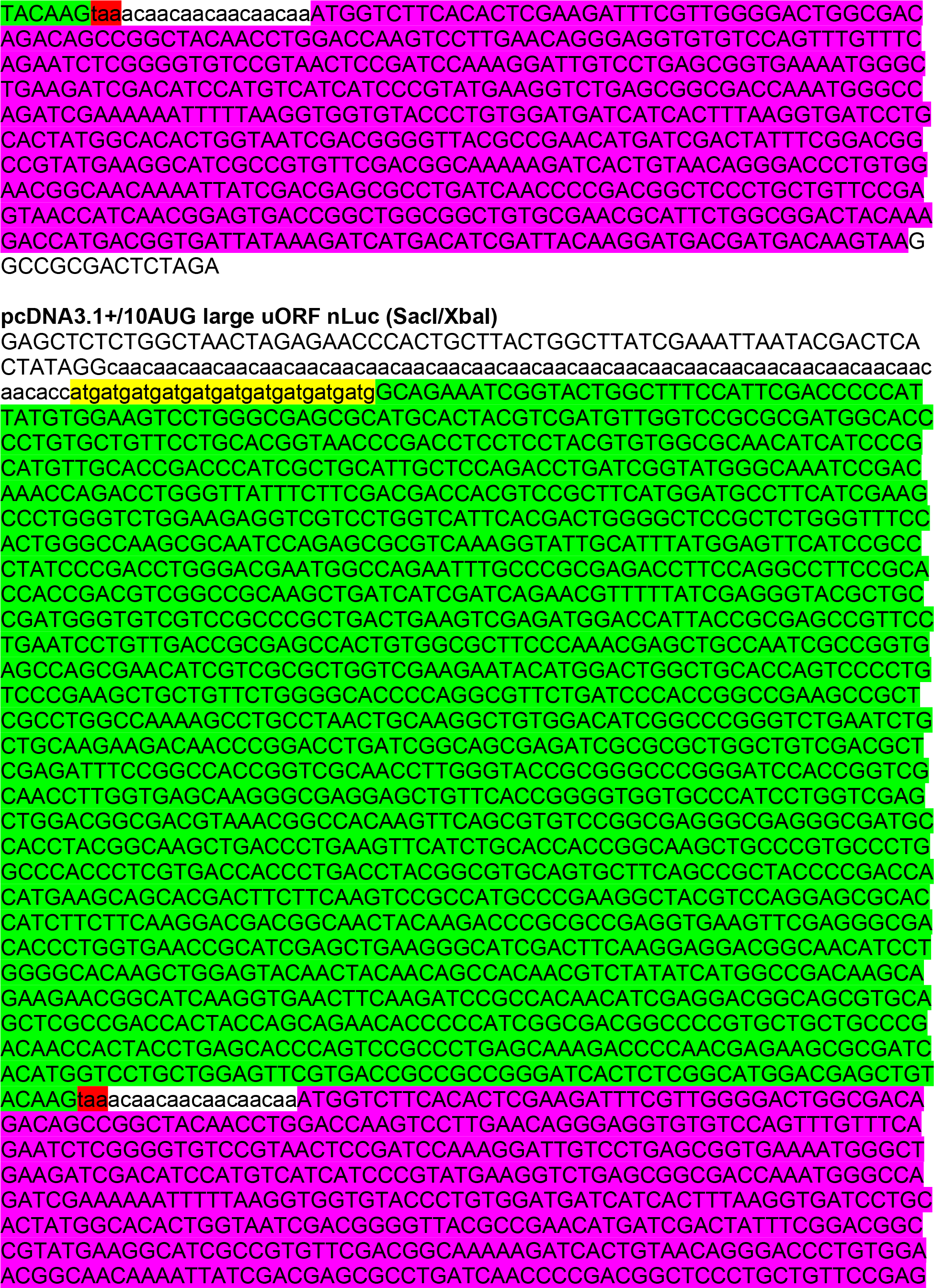
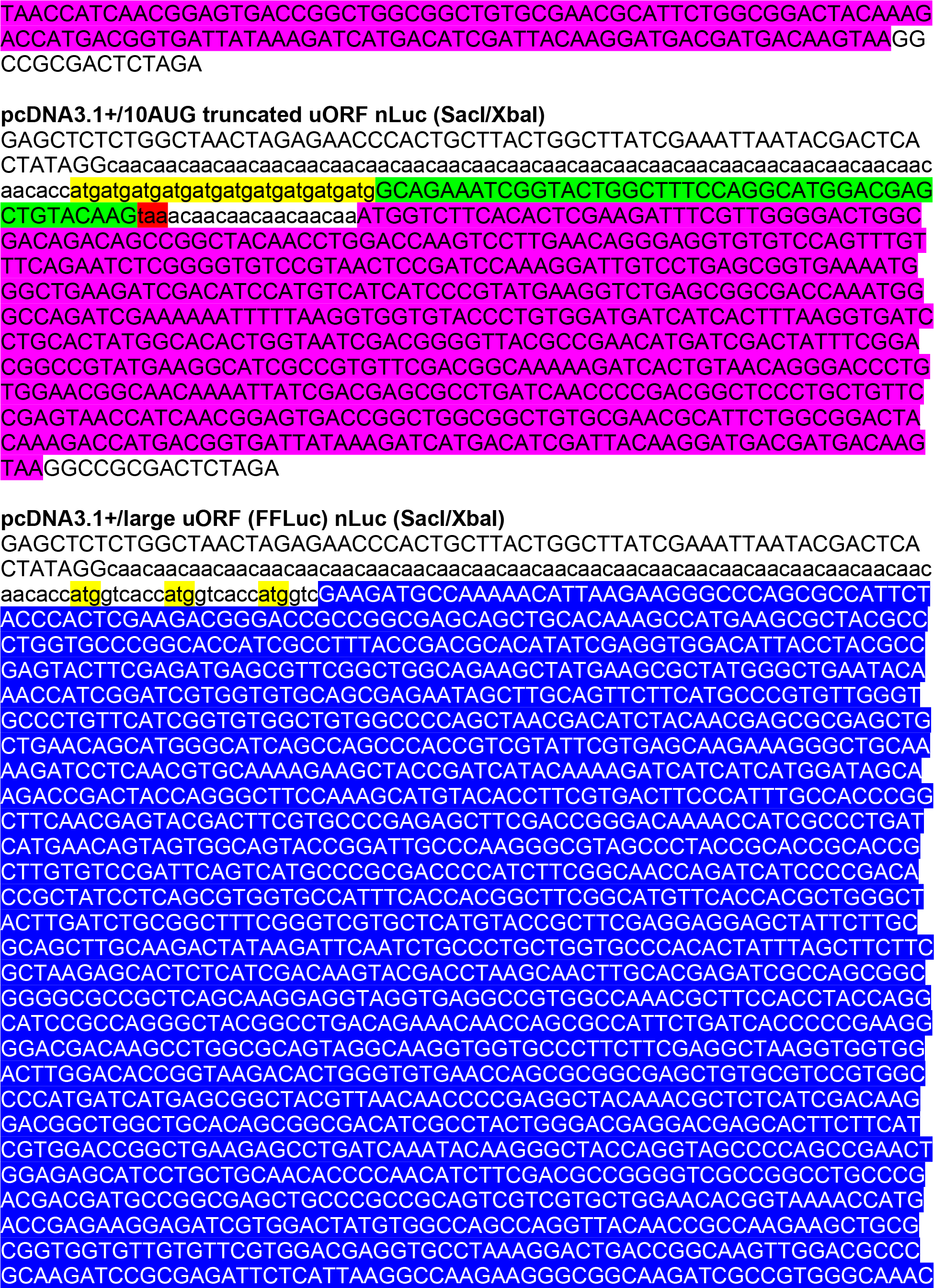
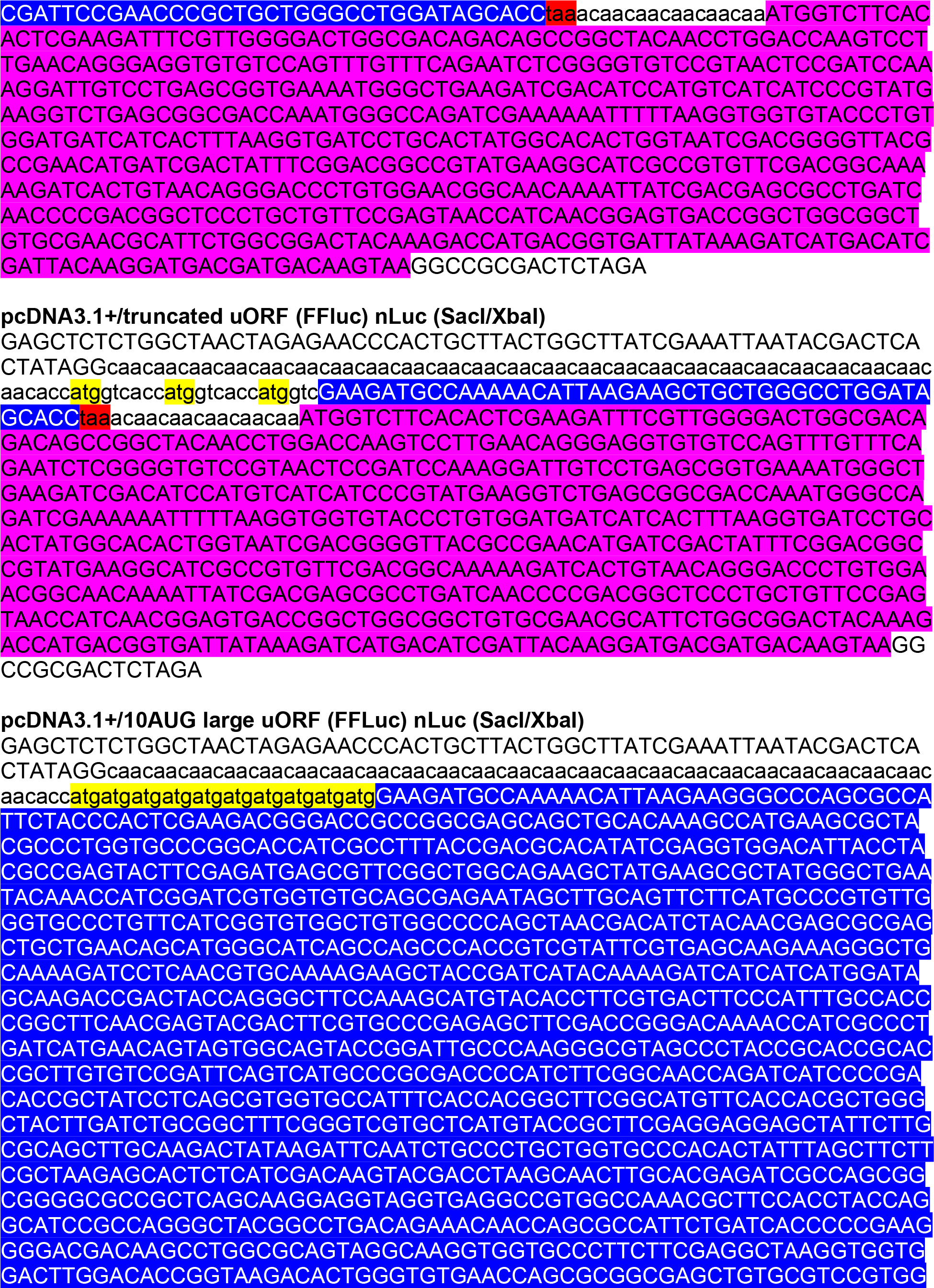
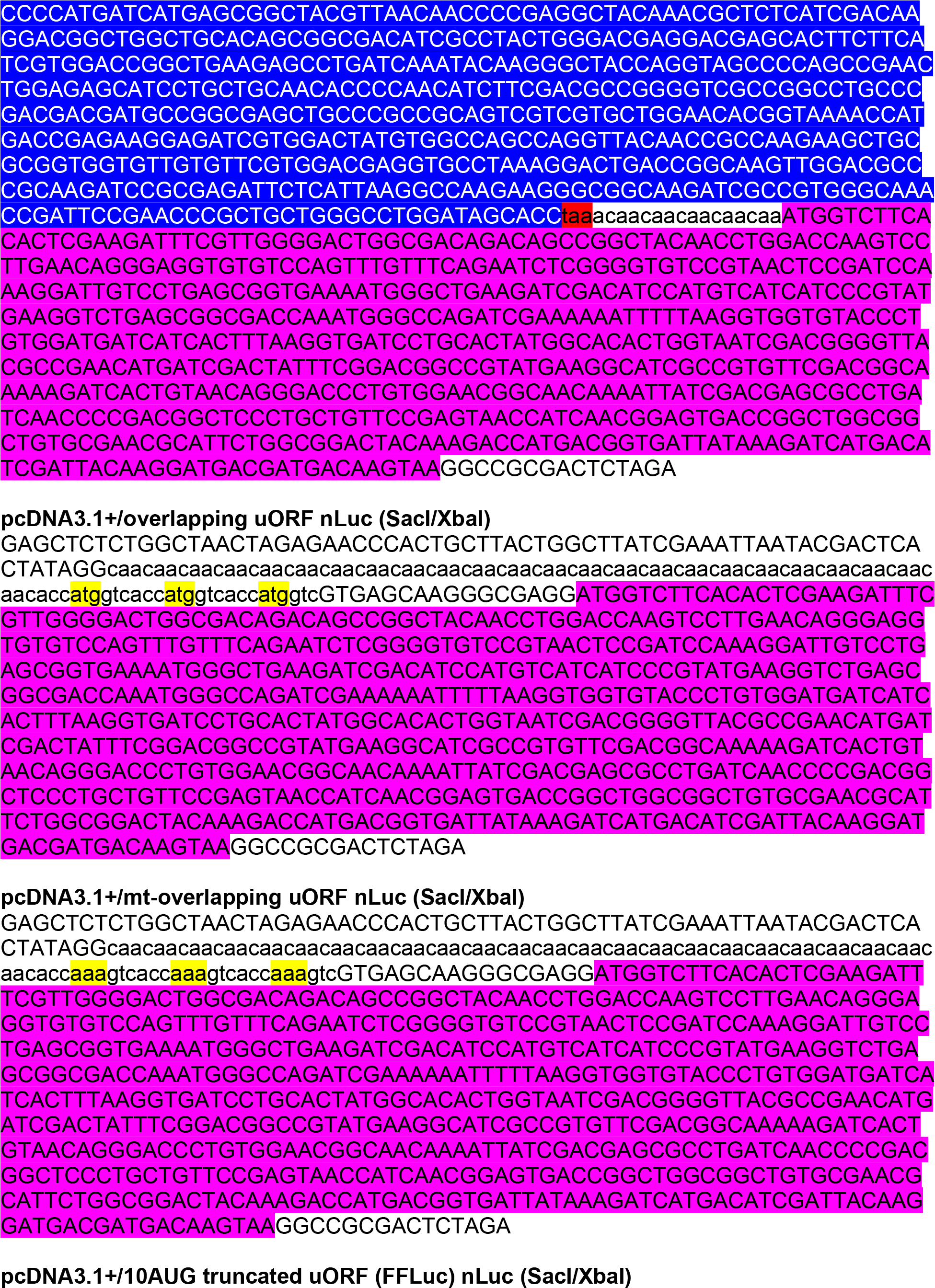
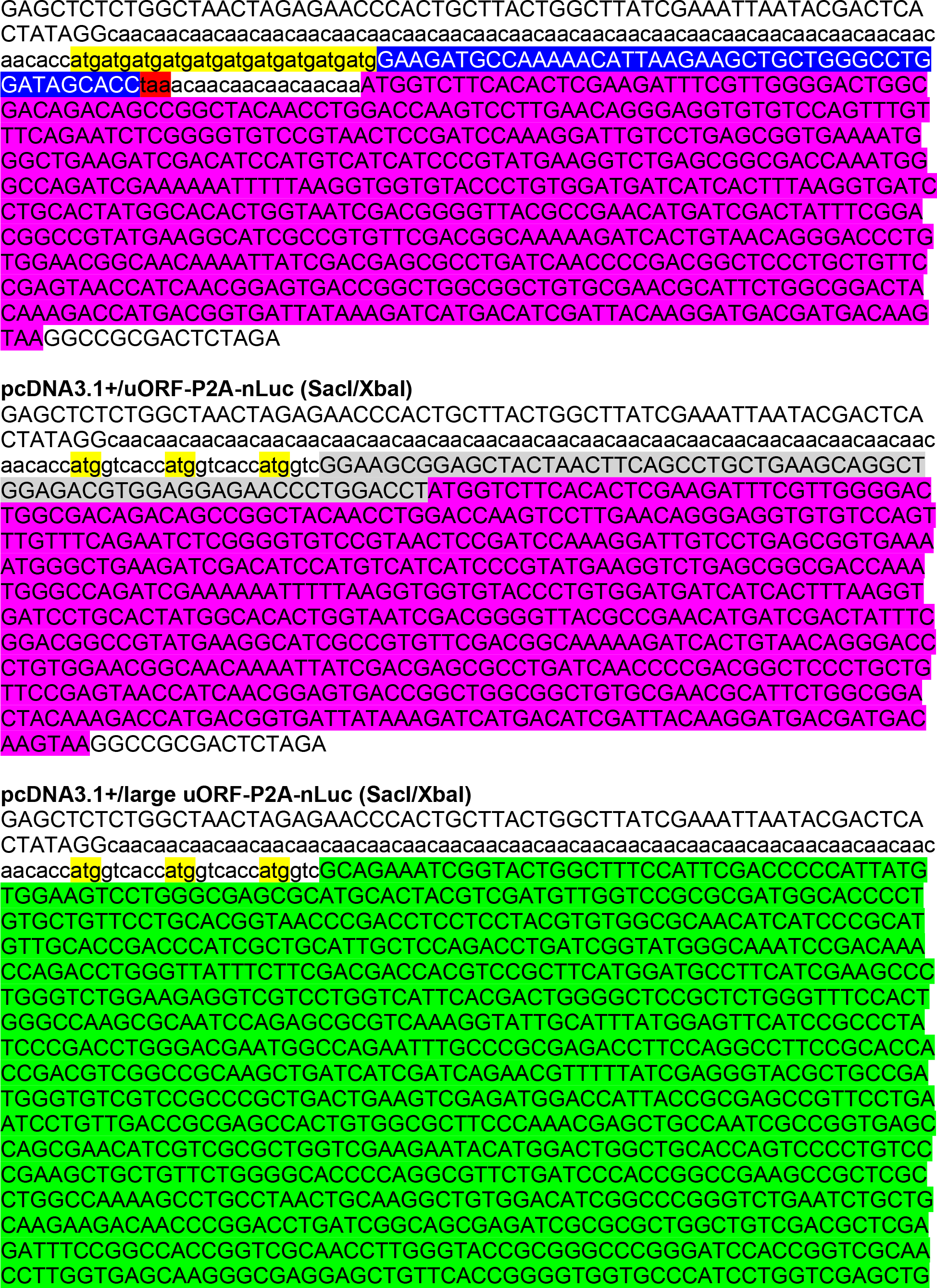
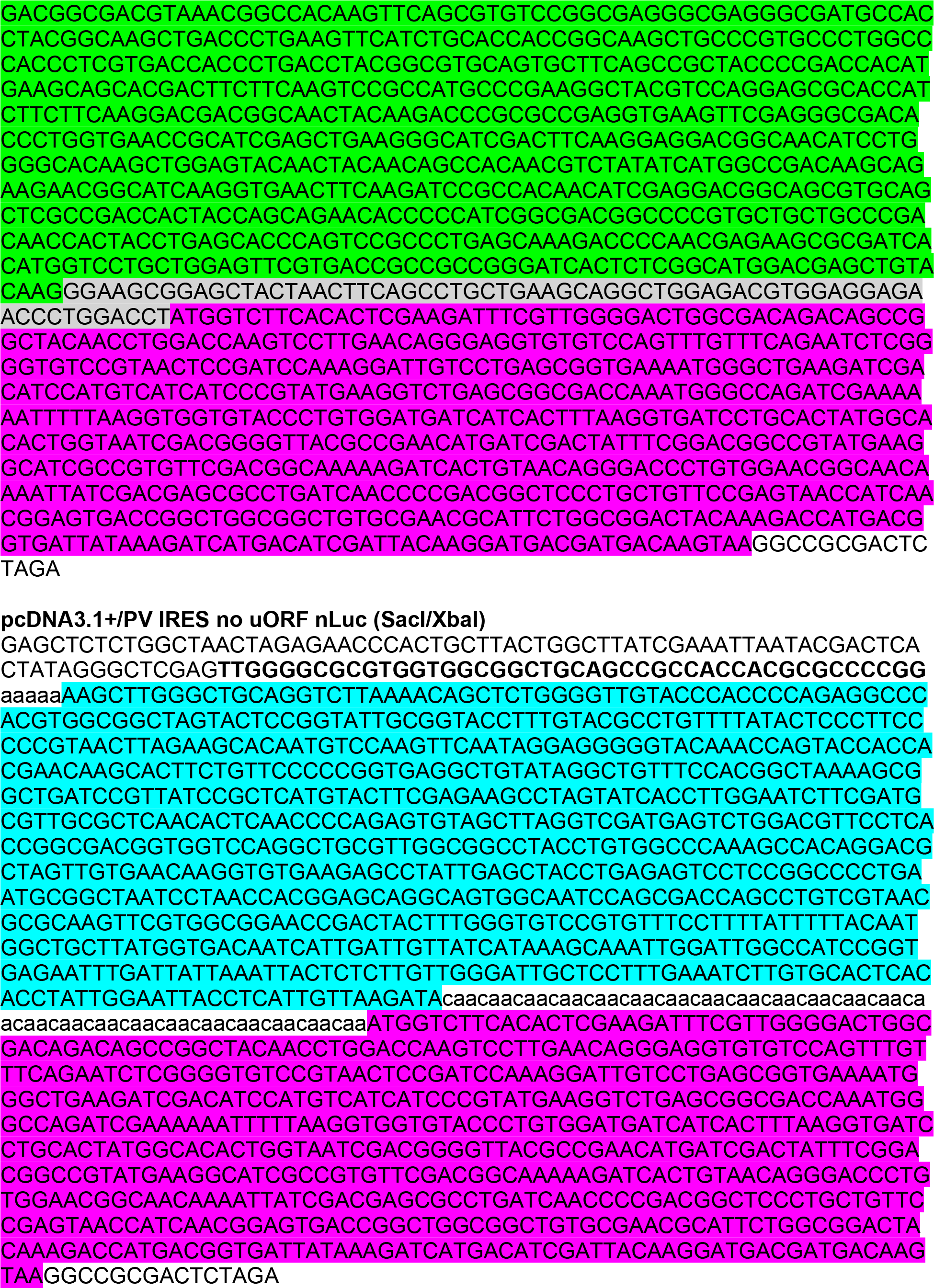
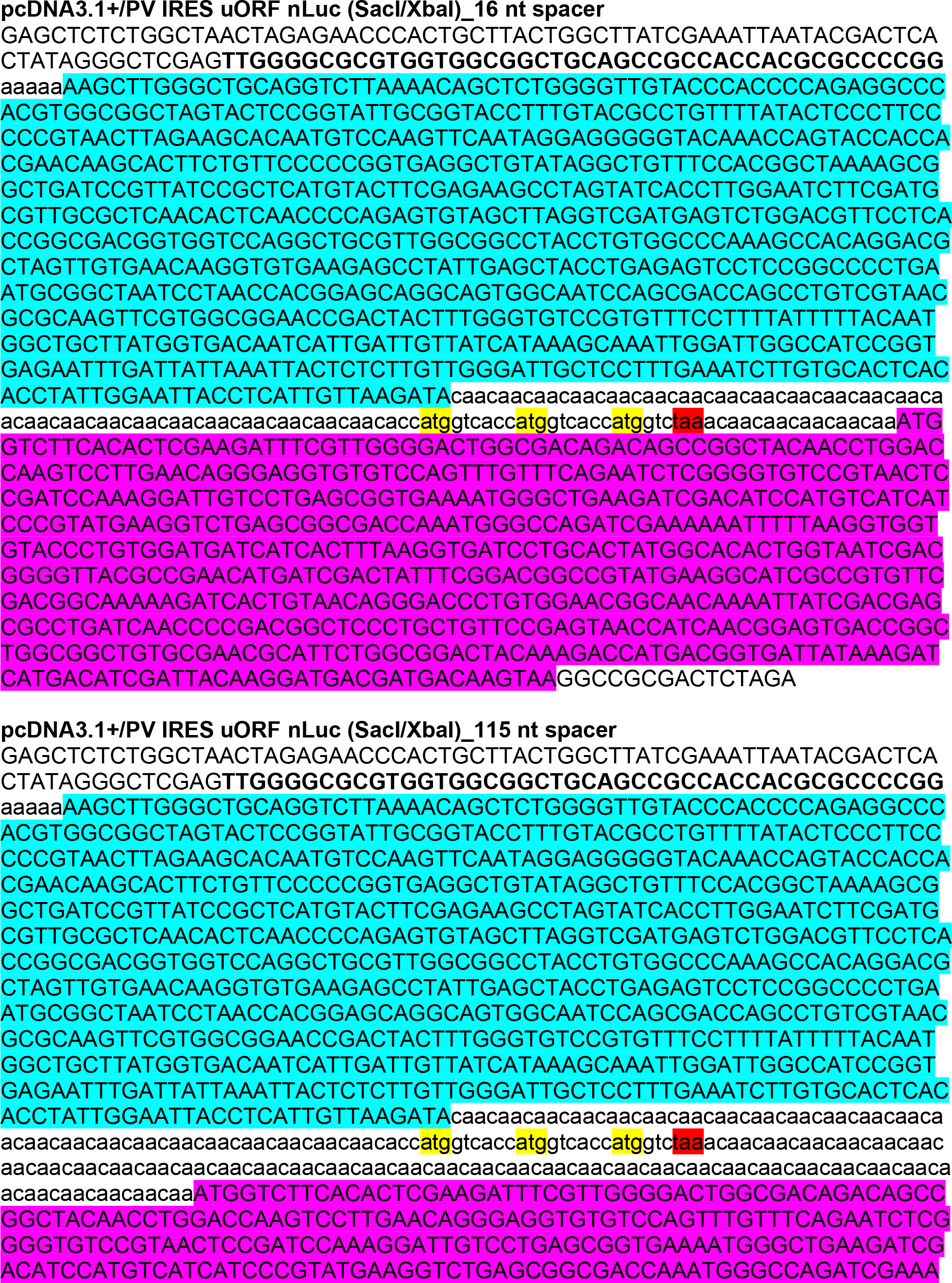
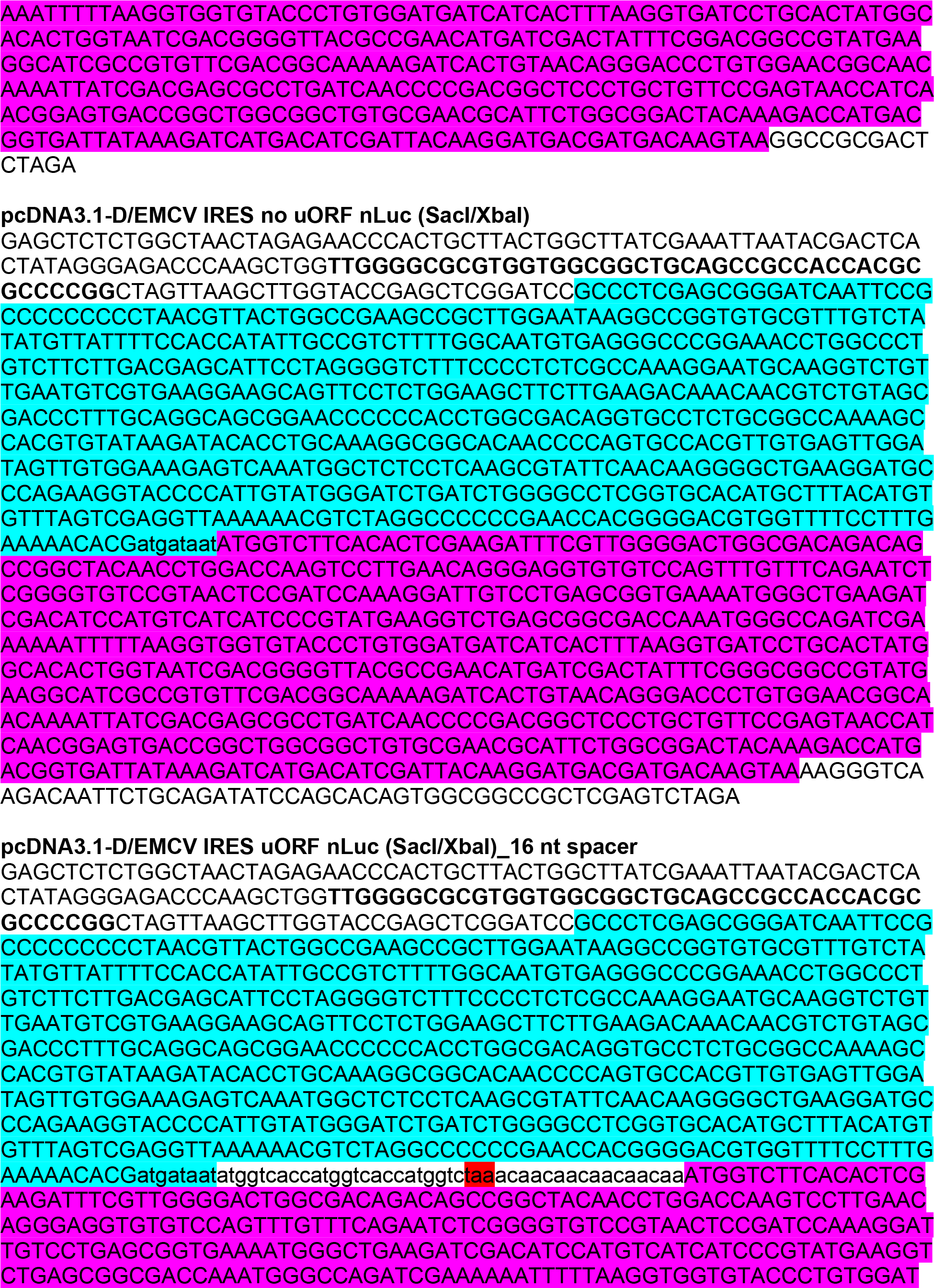
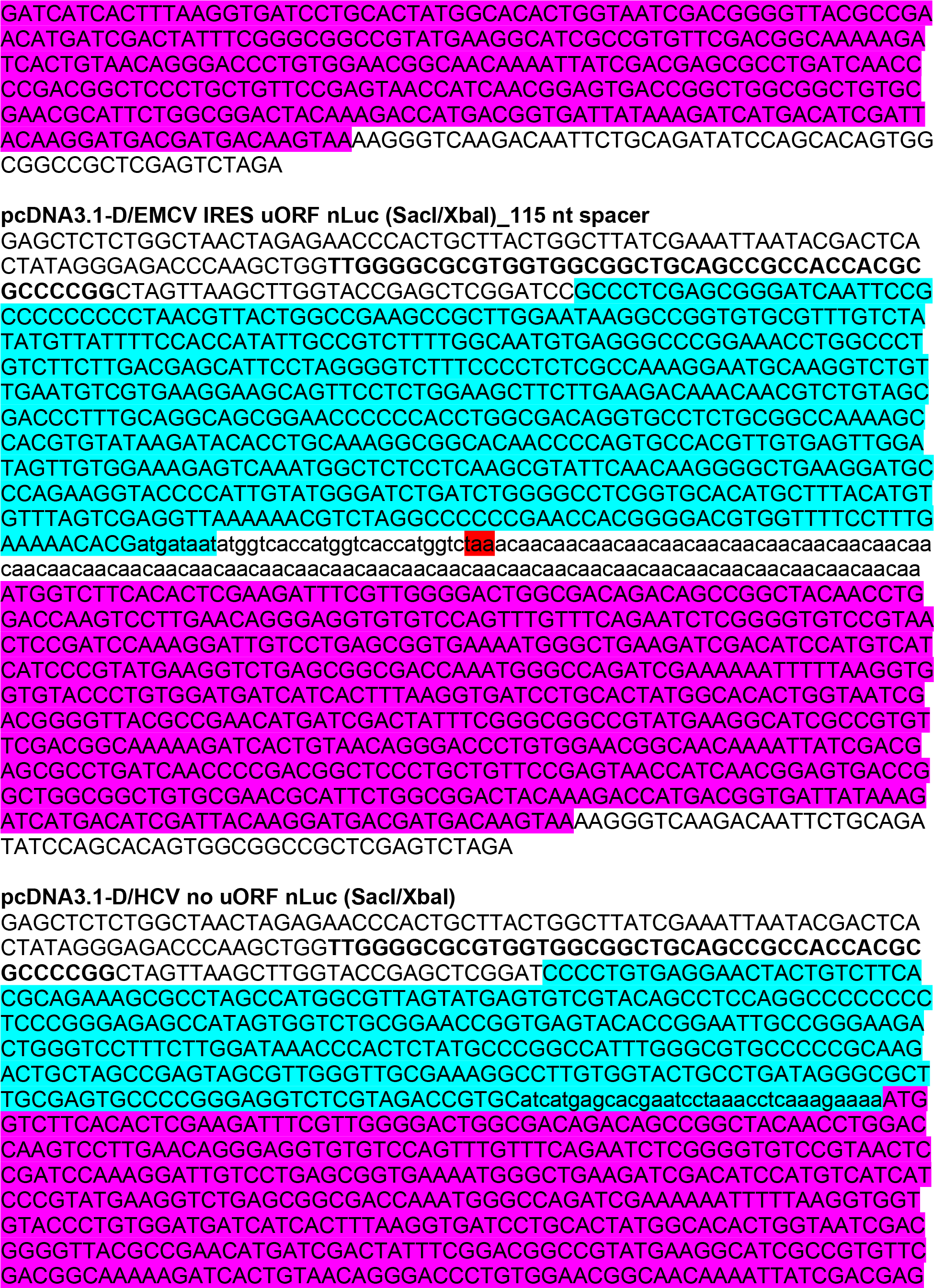
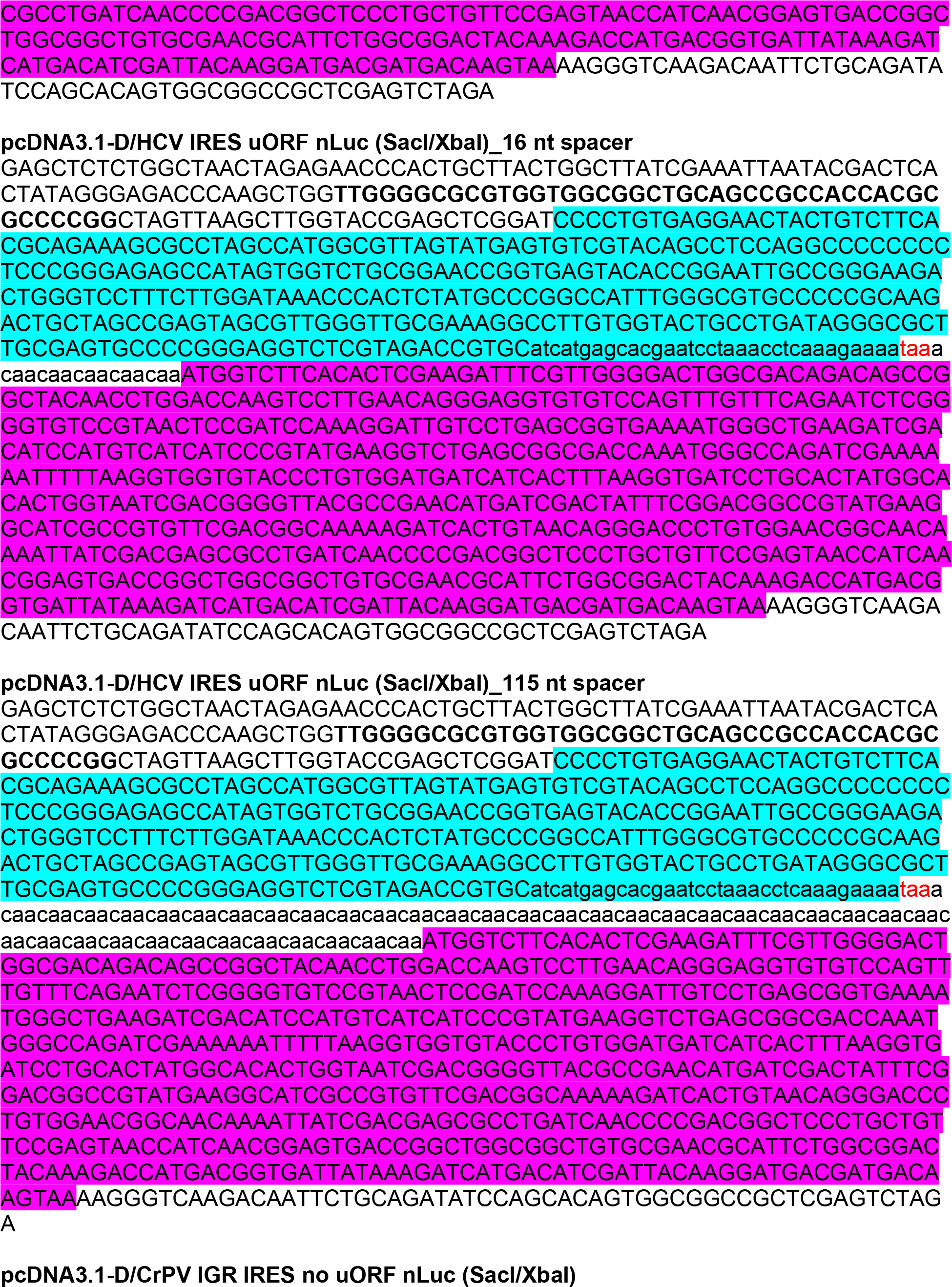
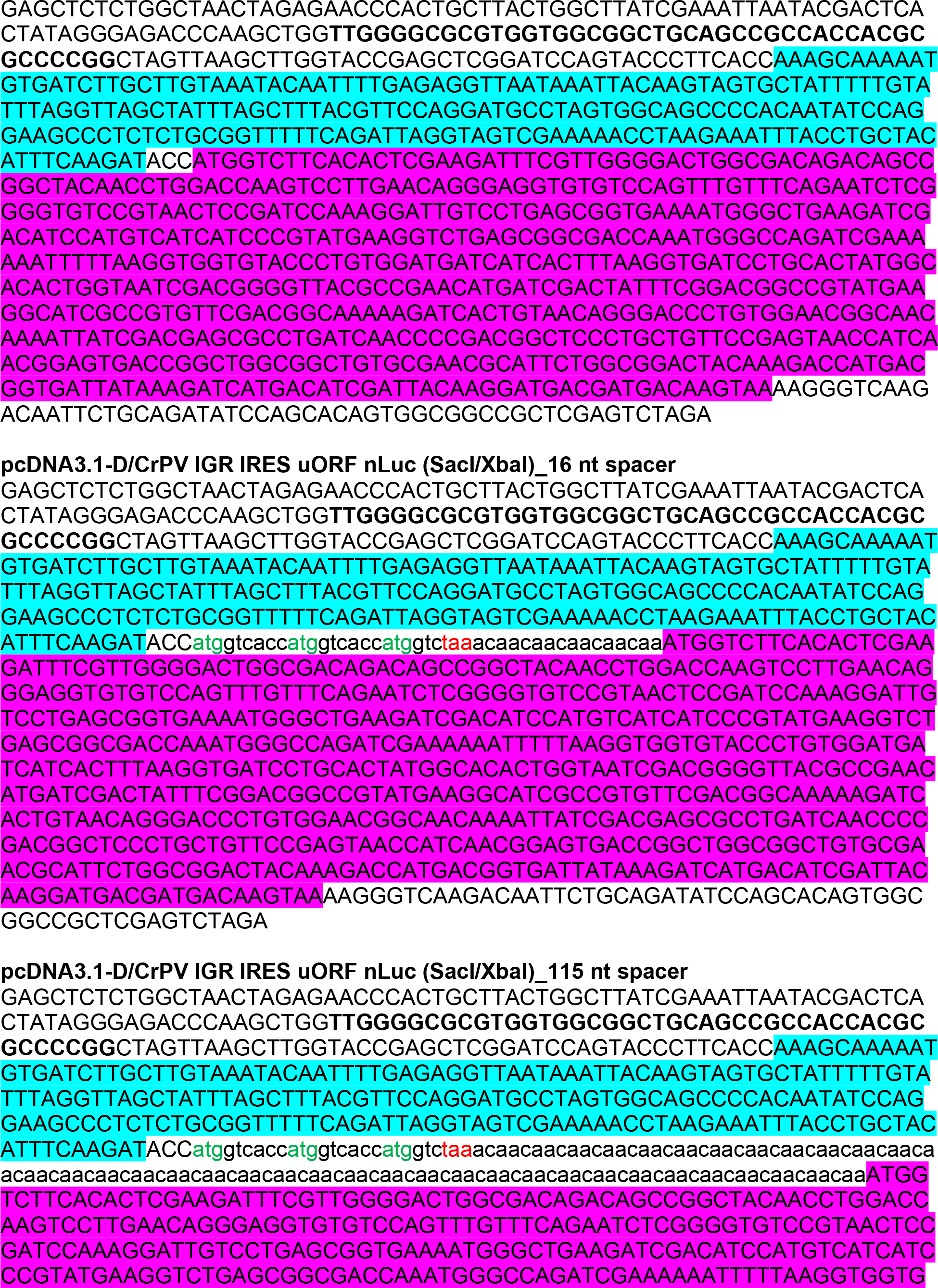
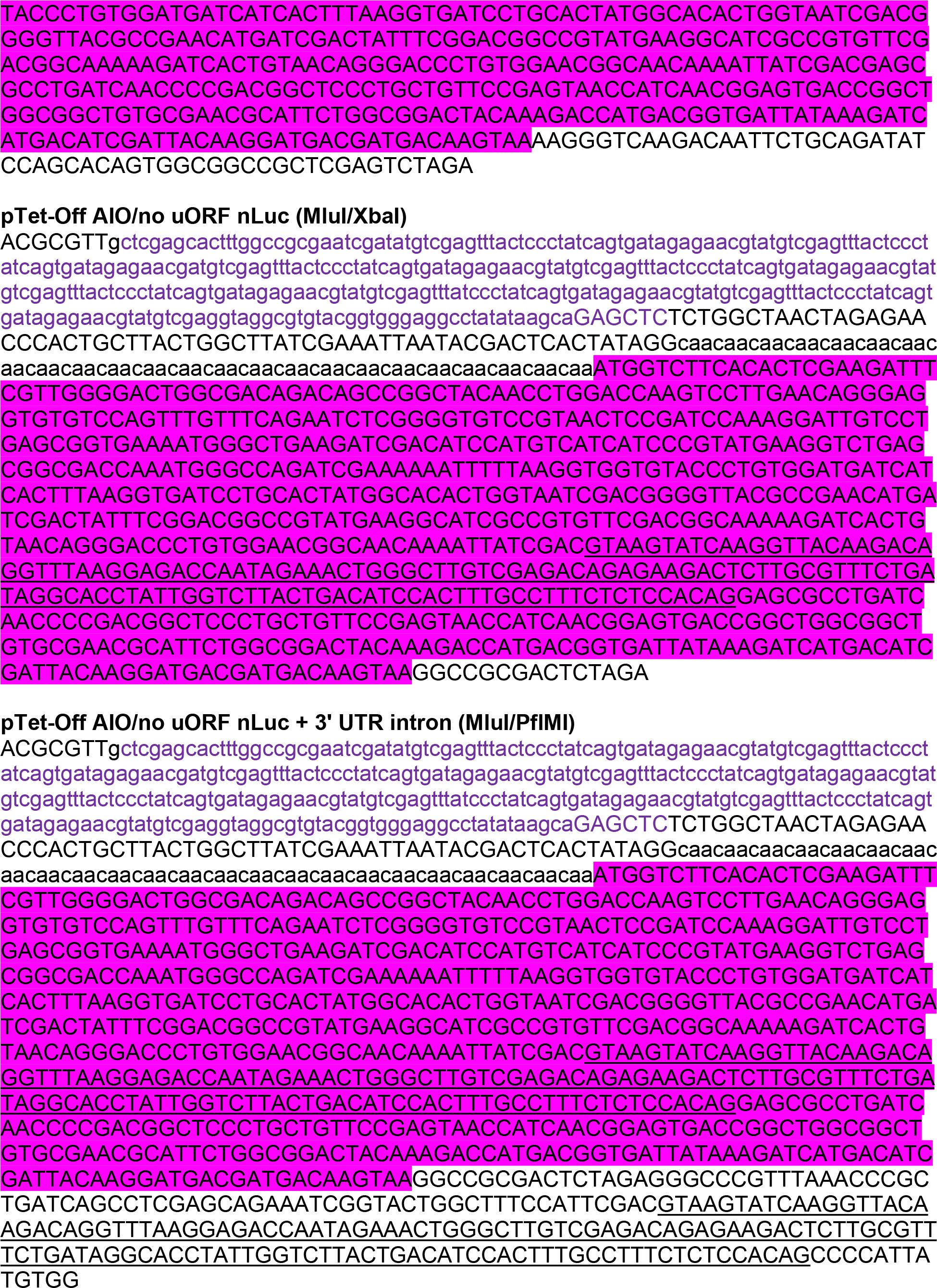
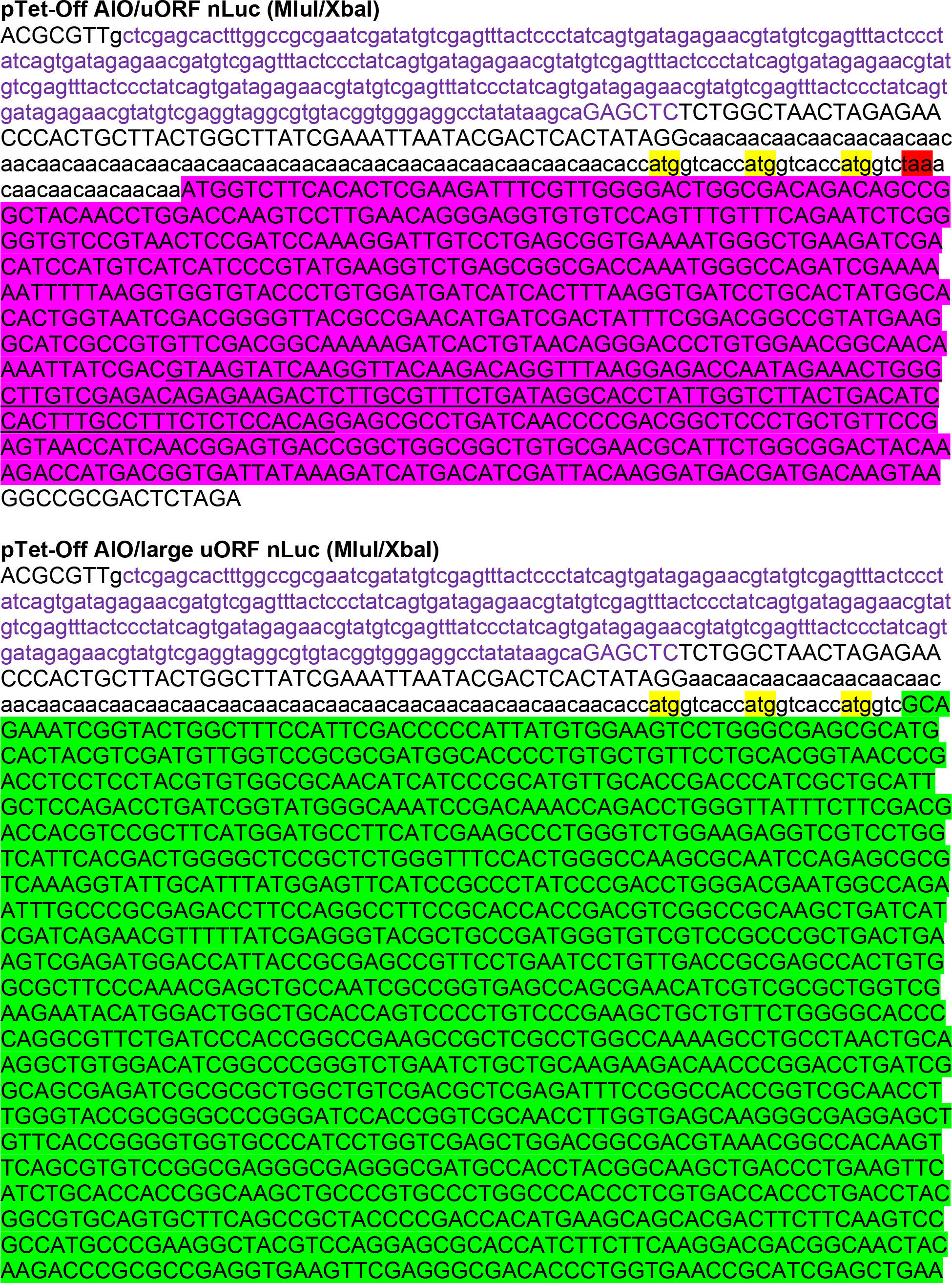
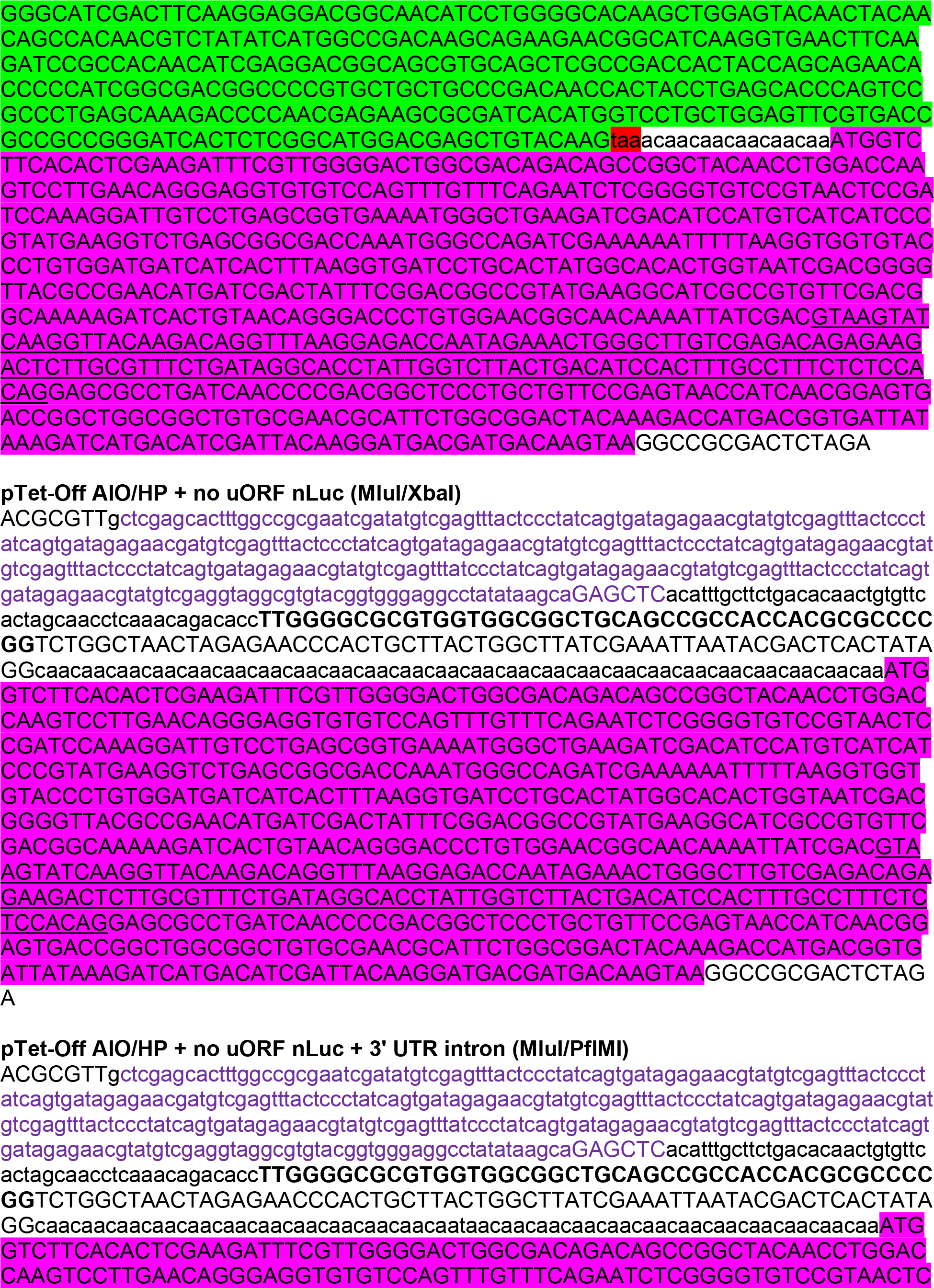
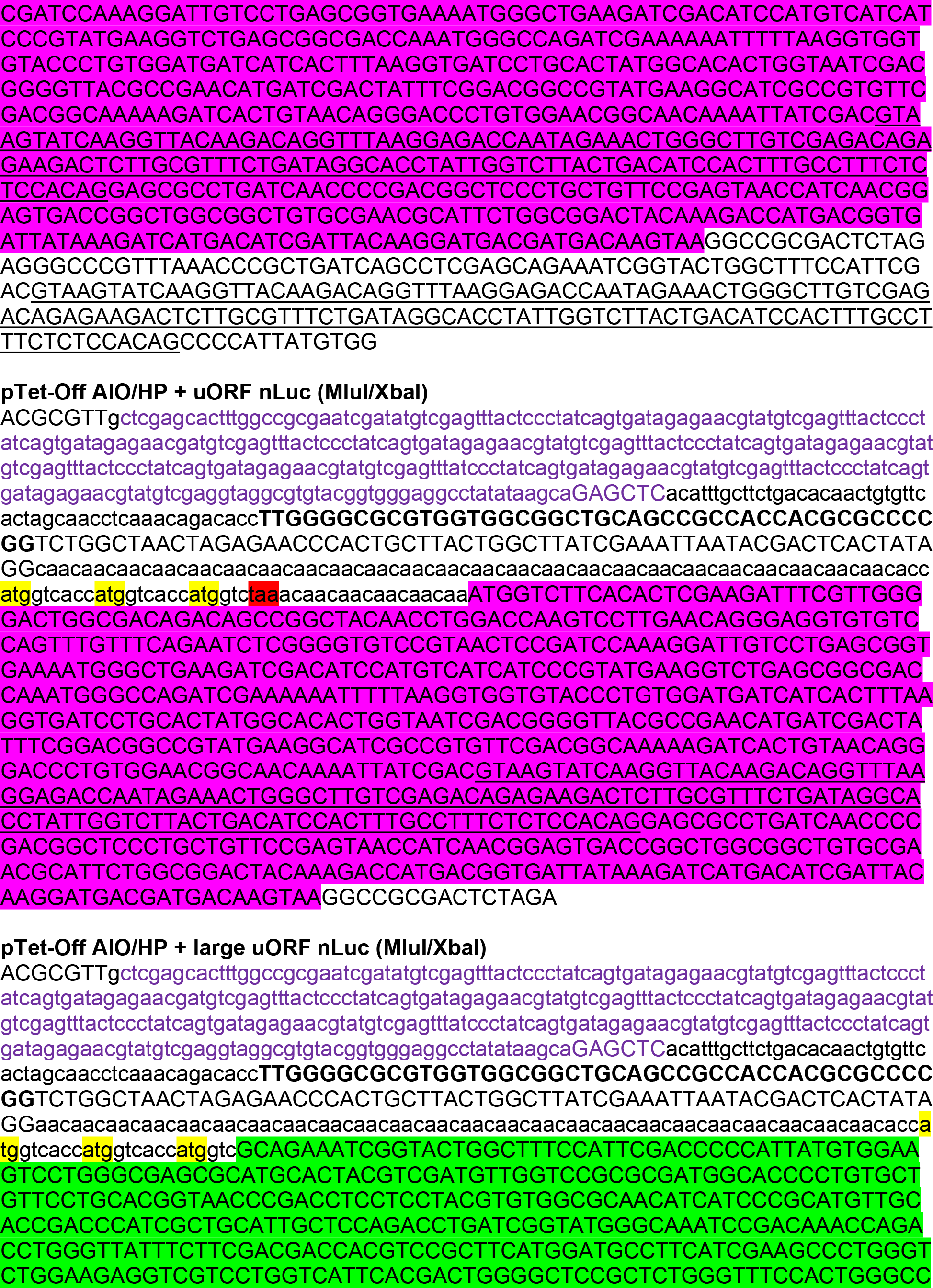
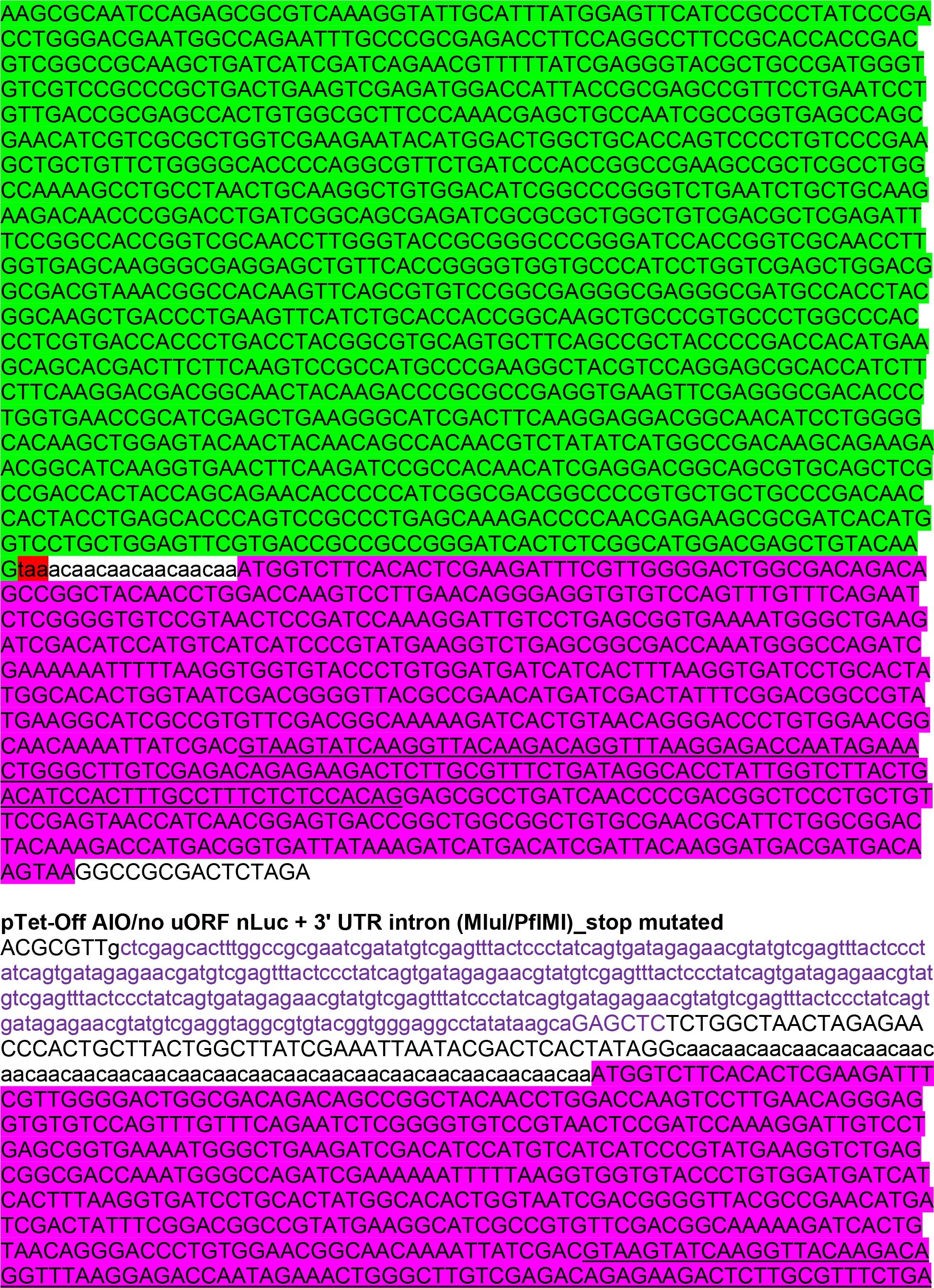
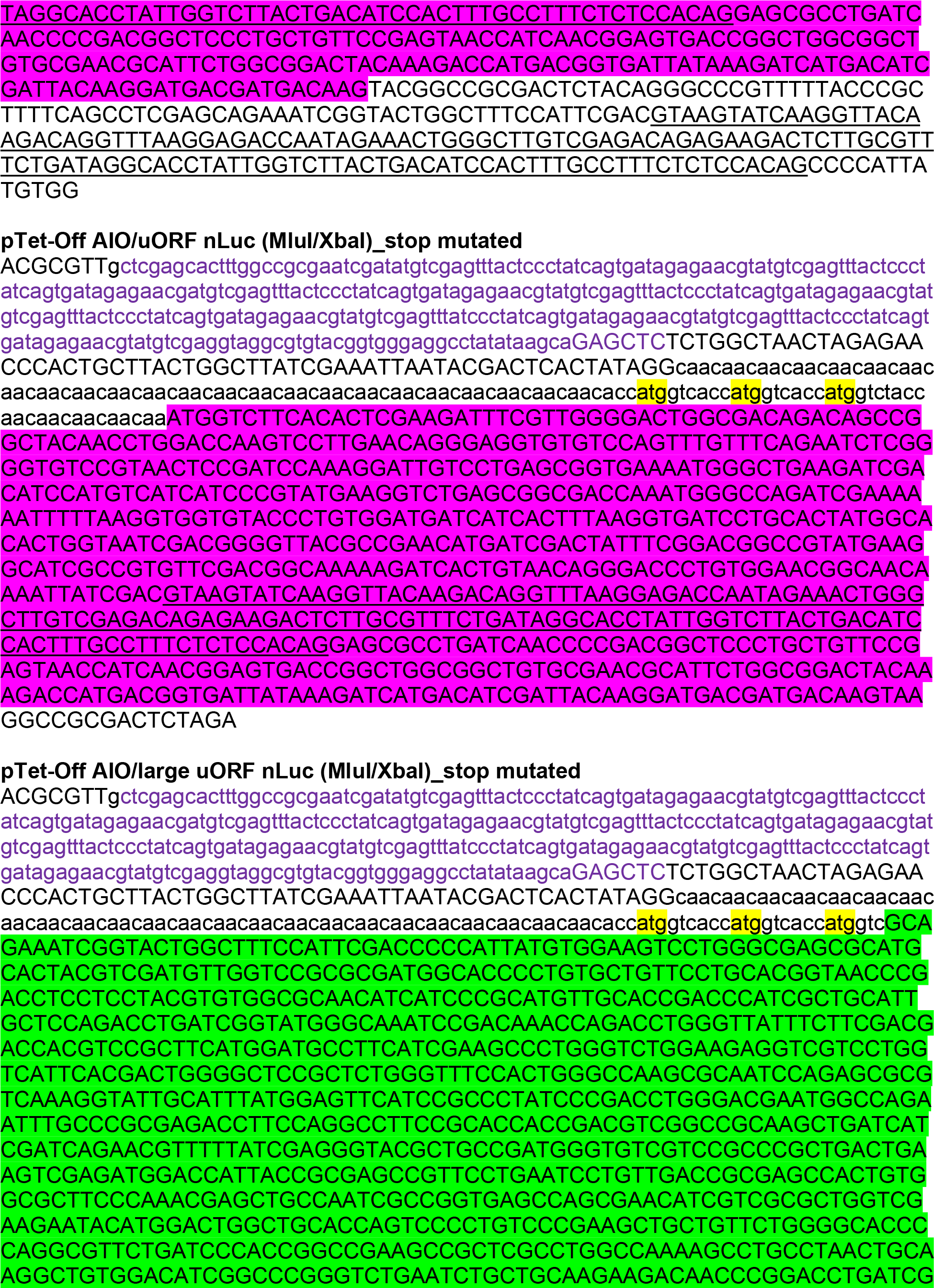
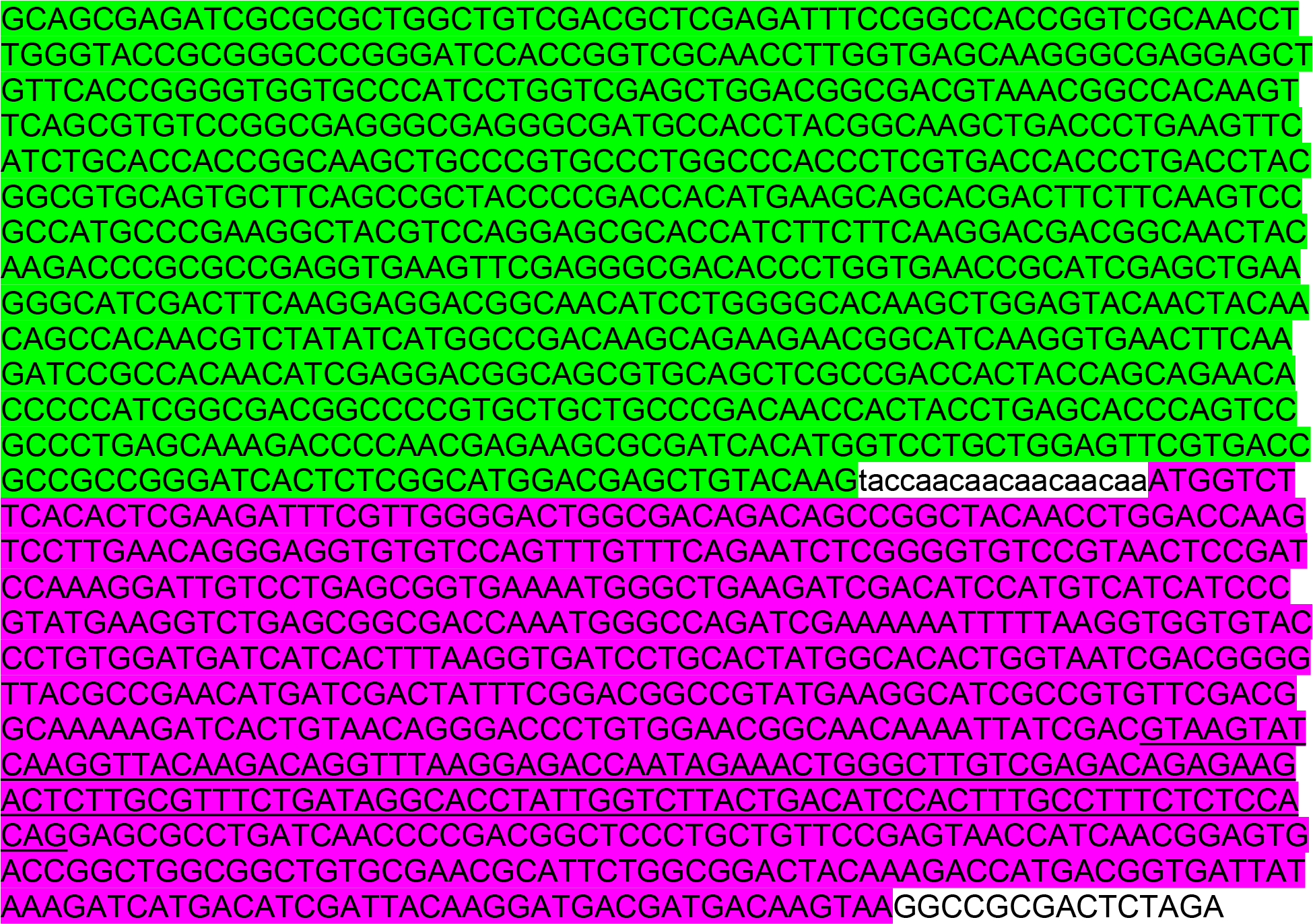

## REFERENCES

1. Annibaldis G, Domanski M, Dreos R, Contu L, Carl S, Klay N, Muhlemann O. 2020. Readthrough of stop codons under limiting ABCE1 concentration involves frameshifting and inhibits nonsense-mediated mRNA decay. Nucleic Acids Res 48: 10259–10279. 10.1093/nar/gkaa758

2. Bohlen J, Fenzl K, Kramer G, Bukau B, Teleman AA. 2020. Selective 40S Footprinting Reveals Cap-Tethered Ribosome Scanning in Human Cells. Mol Cell 79: 561–574 e565. 10.1016/j.molcel.2020.06.005

3. Brito Querido J, Sokabe M, Kraatz S, Gordiyenko Y, Skehel JM, Fraser CS, Ramakrishnan V. 2020. Structure of a human 48S translational initiation complex. Science 369: 1220–1227. 10.1126/science.aba4904

4. Buisson M, Anczukow O, Zetoune AB, Ware MD, Mazoyer S. 2006. The 185delAG mutation (c.68_69delAG) in the BRCA1 gene triggers translation reinitiation at a downstream AUG codon. Hum Mutat 27: 1024–1029. 10.1002/humu.20384

5. Calvo SE, Pagliarini DJ, Mootha VK. 2009. Upstream open reading frames cause widespread reduction of protein expression and are polymorphic among humans. Proc Natl Acad Sci U S A 106: 7507–7512. 10.1073/pnas.0810916106

6. Dyle MC, Kolakada D, Cortazar MA, Jagannathan S. 2020. How to get away with nonsense: Mechanisms and consequences of escape from nonsense-mediated RNA decay. Wiley Interdiscip Rev RNA 11: e1560. 10.1002/wrna.1560

7. Embree CM, Abu-Alhasan R, Singh G. 2022. Features and factors that dictate if terminating ribosomes cause or counteract nonsense-mediated mRNA decay. J Biol Chem 298: 102592. 10.1016/j.jbc.2022.102592

8. Gardner LB. 2008. Hypoxic inhibition of nonsense-mediated RNA decay regulates gene expression and the integrated stress response. Mol Cell Biol 28: 3729–3741. 10.1128/MCB.02284-07

9. Gunisova S, Hronova V, Mohammad MP, Hinnebusch AG, Valasek LS. 2018. Please do not recycle! Translation reinitiation in microbes and higher eukaryotes. FEMS Microbiol Rev 42: 165–192. 10.1093/femsre/fux059

10. Hinnebusch AG. 2017. Structural Insights into the Mechanism of Scanning and Start Codon Recognition in Eukaryotic Translation Initiation. Trends Biochem Sci 42: 589–611. 10.1016/j.tibs.2017.03.004

11. Hulsebos TJ, Kenter S, Verhagen WI, Baas F, Flucke U, Wesseling P. 2014. Premature termination of SMARCB1 translation may be followed by reinitiation in schwannomatosis- associated schwannomas, but results in absence of SMARCB1 expression in rhabdoid tumors. Acta Neuropathol 128: 439–448. 10.1007/s00401-014-1281-3

12. Imamachi N, Salam KA, Suzuki Y, Akimitsu N. 2017. A GC-rich sequence feature in the 3’ UTR directs UPF1-dependent mRNA decay in mammalian cells. Genome Res 27: 407–418. 10.1101/gr.206060.116

13. Jaafar ZA, Oguro A, Nakamura Y, Kieft JS. 2016. Translation initiation by the hepatitis C virus IRES requires eIF1A and ribosomal complex remodeling. Elife 5. 10.7554/eLife.21198

14. Jackson RJ, Hellen CU, Pestova TV. 2010. The mechanism of eukaryotic translation initiation and principles of its regulation. Nat Rev Mol Cell Biol 11: 113–127. 10.1038/nrm2838

15. Jagannathan S, Bradley RK. 2016. Translational plasticity facilitates the accumulation of nonsense genetic variants in the human population. Genome Res 26: 1639–1650. 10.1101/gr.205070.116

16. Karam R, Lou CH, Kroeger H, Huang L, Lin JH, Wilkinson MF. 2015. The unfolded protein response is shaped by the NMD pathway. EMBO Rep 16: 599–609. 10.15252/embr.201439696

17. Kearse MG, Green KM, Krans A, Rodriguez CM, Linsalata AE, Goldstrohm AC, Todd PK. 2016. CGG Repeat-Associated Non-AUG Translation Utilizes a Cap-Dependent Scanning Mechanism of Initiation to Produce Toxic Proteins. Mol Cell 62: 314–322. 10.1016/j.molcel.2016.02.034

18. Kozak M. 1986. Point mutations define a sequence flanking the AUG initiator codon that modulates translation by eukaryotic ribosomes. Cell 44: 283–292. 10.1016/0092-8674(86)90762-2

19. Kozak M. 1987. Effects of intercistronic length on the efficiency of reinitiation by eucaryotic ribosomes. Mol Cell Biol 7: 3438–3445. 10.1128/mcb.7.10.3438-3445.1987

20. Kozak M. 1991. Structural features in eukaryotic mRNAs that modulate the initiation of translation. J Biol Chem 266: 19867–19870.

21. Kozak M. 1997. Recognition of AUG and alternative initiator codons is augmented by G in position +4 but is not generally affected by the nucleotides in positions +5 and +6. EMBO J 16: 2482–2492. 10.1093/emboj/16.9.2482

22. Lapointe CP, Grosely R, Sokabe M, Alvarado C, Wang J, Montabana E, Villa N, Shin BS, Dever TE, Fraser CS, et al. 2022. eIF5B and eIF1A reorient initiator tRNA to allow ribosomal subunit joining. Nature 607: 185–190. 10.1038/s41586-022-04858-z

23. Liu Z, Chen O, Wall JBJ, Zheng M, Zhou Y, Wang L, Vaseghi HR, Qian L, Liu J. 2017. Systematic comparison of 2A peptides for cloning multi-genes in a polycistronic vector. Sci Rep 7: 2193. 10.1038/s41598-017-02460-2

24. Lu PD, Harding HP, Ron D. 2004. Translation reinitiation at alternative open reading frames regulates gene expression in an integrated stress response. J Cell Biol 167: 27–33. 10.1083/jcb.200408003

25. Lykke-Andersen S, Jensen TH. 2015. Nonsense-mediated mRNA decay: an intricate machinery that shapes transcriptomes. Nat Rev Mol Cell Biol 16: 665–677. 10.1038/nrm4063

26. Maekawa S, Imamachi N, Irie T, Tani H, Matsumoto K, Mizutani R, Imamura K, Kakeda M, Yada T, Sugano S, et al. 2015. Analysis of RNA decay factor mediated RNA stability contributions on RNA abundance. BMC Genomics 16: 154. 10.1186/s12864-015-1358-y

27. Martin L, Grigoryan A, Wang D, Wang J, Breda L, Rivella S, Cardozo T, Gardner LB. 2014. Identification and characterization of small molecules that inhibit nonsense-mediated RNA decay and suppress nonsense p53 mutations. Cancer Res 74: 3104–3113. 10.1158/0008-5472.CAN-13-2235

28. Paulsen M, Lund C, Akram Z, Winther JR, Horn N, Moller LB. 2006. Evidence that translation reinitiation leads to a partially functional Menkes protein containing two copper-binding sites. Am J Hum Genet 79: 214–229. 10.1086/505407

29. Pavitt GD, Ron D. 2012. New insights into translational regulation in the endoplasmic reticulum unfolded protein response. Cold Spring Harb Perspect Biol 4. 10.1101/cshperspect.a012278

30. Pereira FJ, Teixeira A, Kong J, Barbosa C, Silva AL, Marques-Ramos A, Liebhaber SA, Romao L. 2015. Resistance of mRNAs with AUG-proximal nonsense mutations to nonsense-mediated decay reflects variables of mRNA structure and translational activity. Nucleic Acids Res 43: 6528–6544. 10.1093/nar/gkv588

31. Pestova TV, Hellen CU. 2003. Translation elongation after assembly of ribosomes on the Cricket paralysis virus internal ribosomal entry site without initiation factors or initiator tRNA. Genes Dev 17: 181–186. 10.1101/gad.1040803

32. Petrov A, Grosely R, Chen J, O’Leary SE, Puglisi JD. 2016. Multiple Parallel Pathways of Translation Initiation on the CrPV IRES. Mol Cell 62: 92–103. 10.1016/j.molcel.2016.03.020

33. Schneider-Poetsch T, Ju J, Eyler DE, Dang Y, Bhat S, Merrick WC, Green R, Shen B, Liu JO. 2010. Inhibition of eukaryotic translation elongation by cycloheximide and lactimidomycin. Nat Chem Biol 6: 209–217. 10.1038/nchembio.304

34. Shirokikh NE, Preiss T. 2018. Translation initiation by cap-dependent ribosome recruitment: Recent insights and open questions. Wiley Interdiscip Rev RNA 9: e1473. 10.1002/wrna.1473

35. Stump MR, Gong Q, Packer JD, Zhou Z. 2012. Early LQT2 nonsense mutation generates N-terminally truncated hERG channels with altered gating properties by the reinitiation of translation. J Mol Cell Cardiol 53: 725–733. 10.1016/j.yjmcc.2012.08.021

36. Szymczak AL, Vignali DA. 2005. Development of 2A peptide-based strategies in the design of multicistronic vectors. Expert Opin Biol Ther 5: 627–638. 10.1517/14712598.5.5.627

37. Tani H, Mizutani R, Salam KA, Tano K, Ijiri K, Wakamatsu A, Isogai T, Suzuki Y, Akimitsu N. 2012. Genome-wide determination of RNA stability reveals hundreds of short-lived noncoding transcripts in mammals. Genome Res 22: 947–956. 10.1101/gr.130559.111

38. Vattem KM, Wek RC. 2004. Reinitiation involving upstream ORFs regulates ATF4 mRNA translation in mammalian cells. Proc Natl Acad Sci U S A 101: 11269–11274. 10.1073/pnas.0400541101

39. Wagner S, Herrmannova A, Hronova V, Gunisova S, Sen ND, Hannan RD, Hinnebusch AG, Shirokikh NE, Preiss T, Valasek LS. 2020. Selective Translation Complex Profiling Reveals Staged Initiation and Co-translational Assembly of Initiation Factor Complexes. Mol Cell 79: 546–560 e547. 10.1016/j.molcel.2020.06.004

40. Wang J, Shin BS, Alvarado C, Kim JR, Bohlen J, Dever TE, Puglisi JD. 2022. Rapid 40S scanning and its regulation by mRNA structure during eukaryotic translation initiation. Cell. 10.1016/j.cell.2022.10.005

41. Wangen JR, Green R. 2020. Stop codon context influences genome-wide stimulation of termination codon readthrough by aminoglycosides. Elife 9. 10.7554/eLife.52611

42. Wek RC. 2018. Role of eIF2alpha Kinases in Translational Control and Adaptation to Cellular Stress. Cold Spring Harb Perspect Biol 10. 10.1101/cshperspect.a032870

43. Yi Z, Sanjeev M, Singh G. 2021. The Branched Nature of the Nonsense-Mediated mRNA Decay Pathway. Trends Genet 37: 143–159. 10.1016/j.tig.2020.08.010

44. Young SK, Wek RC. 2016. Upstream Open Reading Frames Differentially Regulate Gene-specific Translation in the Integrated Stress Response. J Biol Chem 291: 16927–16935. 10.1074/jbc.R116.733899

